# Drop on fixed target reaction initiation approach for serial and time resolved crystallography

**DOI:** 10.64898/2025.12.17.694847

**Authors:** Jos J. A. G. Kamps, Philip Hinchliffe, Johan Glerup, Emily I. Freeman, Pauline A. Lang, Catherine L. Tooke, Michael Beer, Laura Parkinson, Do-Heon Gu, Sehan Park, Nicholas Devenish, Tiankun Zhou, Anastasya Shilova, Patrick Rabe, Christopher J. Schofield, James Spencer, Jaehyun Park, Robin L. Owen, Allen M. Orville, Pierre Aller

## Abstract

We describe the design and implementation of a drop on fixed target method for time-resolved serial crystallography at both synchrotron and XFEL facilities. A piezoelectric droplet dispensing pipette is employed for addition of picolitre volume (40 – 90 pL) aqueous droplets, containing (co-)substrate(s) or ligand(s), onto enzyme microcrystals immobilised on a solid support. The system was tested with various enzyme systems, including lysozyme and two ꞵ-lactamases, CTX-M-15 and AmpC_EC_. Mitigation strategies for cross-well contamination, including the implementation of interleaved controls, are described; the overall performance of the system at synchrotron and X-ray free electron laser facilities was evaluated. This drop on fixed target method is a reliable framework for time-resolved crystallography and will improve the consistency of measurements across facilities.

## 1. Introduction

Serial femtosecond crystallography (SFX) on protein crystals became possible with the introduction of hard X-ray free electron laser (XFEL) facilities at the beginning of the 2010s (Emma *et al*., 2010). The “diffraction before destruction” principle (Neutze *et al*., 2000) enables high-resolution structural data collection without radiation-induced modifications (sometimes referred to as “radiation damage”), which is crucial for studying protein crystals with sensitive chemical entities, including disulphides, carboxylic acids, and redox-active metals (Garman & Weik, 2023). The femtosecond (fs) XFEL pulse duration provides an opportunity to probe extremely rapid chemical phenomena, *e.g.* bond isomerisation, in photoactive proteins (Poddar *et al*., 2022). The destructive nature of the XFEL pulse, however, necessitates a continuous feed of fresh crystalline sample to be supplied for SFX data collection. In practice, this requirement places substantial demands on sample preparation and efficient use of facility time, because XFEL beamtime remains scarce due to its inherent design and limited number of operational facilities. The serial crystallography approach has also been adopted and developed at microfocus beamlines at synchrotrons (serial synchrotron crystallography, SSX), offering low dose room temperature data collection, while improving accessibility (Pearson *et al*., 2020).

Early SFX studies often focused on enzymes available in large quantities, due to the high sample consumption rates (10 – 40 µL min^-1^) required by sample delivery systems using gas dynamic virtual nozzles (GDVNs) (Weierstall *et al*., 2012). Efforts to reduce sample consumption for GDVNs include use of a high repetition source pulse and segmented flow sample delivery (Echelmeier *et al*., 2020). High-viscosity extruders (HVEs), which suspend microcrystals in a viscous carrier matrix, substantially reduce the sample consumption rate (10 – 300 nL min^-1^) (Weierstall *et al*., 2014). Voltage-driven electrokinetic inject systems, such as Concentric Microfluidic Electric Sample Holders (CoMESHs), operate at a favourably low flow rate (0.14 – 3.1 µL min^-1^) and use a sister liquid supplemented with cryoprotectant as an outer sheet to prevent freezing (Sierra *et al*., 2016). Hybrid microfluidic methods manifest remarkable sample efficiency (Nam & Cho, 2021; Gu *et al*., 2023). On-demand sample delivery reduces sample consumption by presenting the microcrystal slurry to the X-ray interaction region at the repetition rate of the X-ray source. The drop on demand tape drive setup (Fuller *et al*., 2017; Butryn *et al*., 2021; Kamps *et al*., 2024) uses a conveyor belt system to carry nL-sized droplets (overall rate 3.3 – 10 µL min^-1^) through a reaction initiation region, before presenting the droplets to the X-ray interaction region. The reaction region supports different initiation strategies including gaseous diffusion, drop on drop mixing, or light activation, increasing the number of systems suitable for time-resolved (tr) SSX and SFX.

GVDNs, HVEs, CoMESHs and tape drive systems have demonstrated utility in liquid mixing for small molecule (co-)substrate/ligand applications. For mixing injectors, the accessible mixing time frames for these methods are relatively short, ranging from ms to a few seconds, since the mixing times are closely tied to flow rate (Calvey *et al*., 2019; Pandey *et al*., 2021; Dasgupta *et al*., 2019; Kupitz *et al*., 2017; Olmos *et al*., 2018; Knoška *et al*., 2020). In contrast, HVE-based approaches are better suited for longer mixing times (from 2 to 20 s) (Vakili *et al*., 2023), limiting the systems that can be studied, as typical turnover times average ∼60 ms (Aller & Orville, 2021). Tape-drive based liquid mixing has been implemented using T-junction capillary mixing prior to deposition on the tape, with mixing times ranging from 500 ms to several minutes (Zielinski *et al*., 2022; Beyerlein *et al*., 2017). Alternatively, mixing in a drop-on-drop tape-drive system can be achieved through piezoelectric ejection, where small-molecule-containing droplets are deposited onto microcrystal-containing droplets in an on-demand drop-on-drop approach, inducing turbulent mixing offering mixing time from 10s ms to few seconds (Butryn *et al*., 2021, Nguyen *et al*., 2023).

Microfluidics has been employed at synchrotrons to enhance time-resolved mixing experiments. 3D-MiXD (Monteiro *et al*., 2020) utilizes microfluidic chips that mix microcrystals and ligands before exposing them to X-rays through an open central channel. The time delays for 3D-MiXD range from hundreds of milliseconds to 2 seconds.

Fixed target approaches differ from the aforementioned flowing sample delivery methods, which usually rely on transporting microcrystalline slurry via a capillary system, and instead rely on sandwiching the slurry between two X-ray transparent sheets held within a frame (Sherrell *et al*., 2015, Narayanasamy *et al*., 2025, Jaho *et al*., 2024, Rabe *et al*., 2020). This removes constraints on upper crystal size and avoids challenges of droplet formation or jetting stability. In principle, this enables multiple data collections from the same microcrystal sample. The fixed target approach is highly sample efficient and simple in design. Crystals can be grown directly on the solid support, eliminating the need for mechanical handling for especially delicate samples (Apel *et al*., 2019). The widespread adoption of fixed target systems has encouraged discussions on standardising design and nomenclature to facilitate cross facility exchange (Owen *et al*., 2023).

Fixed target sample delivery systems can be divided into two types: (i) the directed raster type, and (ii) the aperture aligned type (Owen *et al*., 2023). Directed raster systems use a thin layer of microcrystals randomly distributed across an active area between two transparent polymer films. This inexpensive design uses a drawn grid of probing points across the active area for data acquisition. Practical challenges, including uniform distribution of microcrystals, avoidance of wrinkle formation in the two polymer sheets, and preventing crystal settling once the sample is mounted on the beamline, remain and should be accounted for by the user for reliable implementation (Doak *et al*., 2024, 2018). Additionally, grid parameters (*e.g.* the distance between probing spots), must be carefully tuned to avoid overlap between regions affected by diffused radicals generated in previously exposed areas. As such, raster systems offer a robust platform to explore the viability of a novel enzyme system for SSX and SFX experiments and hold potential for future automation. The enclosed design and lack of compartmentalisation, however, constrain their applications in time-resolved experiments. Aperture aligned systems position crystals at predictable locations within funnel-shaped cavities (or wells) that allow liquid to go through and capture crystals of a particular minimum size (Carrillo *et al*., 2023). The microcrystal slurry is loaded onto a solid support (silicon or a microstructured polymer), either manually or using droplet ejection (Davy *et al*., 2019), after which excess mother liquid is removed through blotting or application of vacuum. The crystals settle inside the wells, placing them at predictable locations, and in theory, isolating each well from the neighbouring one.

It is then possible to perform pump-probe tr-SFX and tr-SSX studies using a well-focused and well-aligned laser light (Smyth *et al*., 2025). However, light contamination of adjacent wells is possible, impacting the accuracy of time-resolved experiments (Gotthard *et al*., 2024). This approach could be deployed to investigate light sensitive proteins (Schulz *et al*., 2018), and can, under appropriate conditions, be used to release small molecules such as O_2_ or NO from photolabile precursors (Sandelin *et al*., 2024; Smyth *et al*., 2025). The method could be extended to photoswitchable chemicals, which can be altered between active and inactive states under specific wavelength irradiation (Nasrallah *et al*., 2021).

Recently, Mehrabi and co-workers pioneered the elegant use of droplet addition in combination with a fixed target approach for time-resolved mixing experiments (Mehrabi, Schulz, Agthe *et al*., 2019; Mehrabi, Schulz, Dsouza *et al*., 2019, Schulz *et al*., 2025). Their method uses the aperture-aligned approach to separate microcrystals into individual wells, within each of which reactions are initiated by adding a small molecule containing droplet (∼75 pL; here after referred to as “droplet(s)”). After a time delay ranging from ms to seconds, or longer, the wells are probed with X-rays, enabling time-resolved crystallographic experiments based upon adding nearly any type of ligand(s). Theoretical work has suggested diffusion could be a limiting factor for achieving low-ms mixing times (Schmidt, 2020), even when technical challenges related to sample delivery are overcome. This method expands the use of fixed target approaches for time-resolved macromolecular crystallography (tr-MX).

Here, we describe an adaptation of the drop on fixed target system that uses a piezoelectric droplet injector for the addition of small molecule ligands or substrates as a means for tr-MX at microfocus beamline I24 (Diamond Light Source (DLS), UK) and deployment at the PAL-XFEL (Pohang, Republic of Korea) facility. In particular, we address challenges including cross-well contamination, provide a detailed breakdown of design choices, and present troubleshooting guidelines to help mitigate these challenges. As proof of principle of the drop on fixed target approach, we demonstrate binding of small molecule inhibitors to several enzyme systems: *N*-acetyl-D-glucosamine (GlcNAc) to hen egg white lysozyme (HEWL), avibactam to the class A serine β-lactamase cefotaximase-Munich-15 (CTX-M-15) from *Enterobacterales*, and avibactam to the class C serine β-lactamase AmpC from *Escherichia coli* (AmpC_EC_). Our results demonstrate viability of the drop on fixed target method mixing of small molecule ligands, and show that inclusion of interleaved controls is important for time-point validation.

## 2. Methods

### 2.1 Sample loading

Protein microcrystalline samples were prepared as detailed in the SI.

An adapted procedure (Horrell *et al*., 2021) was used to prepare microcrystalline slurry on silicon chips. Thus, in a controlled relative-humidity (> 80%) enclosure, a microcrystal slurry (30-100 µL) was added to a glow-discharged silicon chip (10 or 15 µm aperture size) immediately prior to use. Excess mother liquor was removed using a gentle vacuum and careful tissue blotting, removing surplus liquid while avoiding crystal dehydrating; this vital step remains a trial-and-error process. Crystals larger than the well aperture remain trapped (Figure S1). The average number of crystals per well depends on the crystal slurry density (≈10^7^–10^8^ crystals/mL) and loaded volume. For resting-state structures, the chip is sandwiched between two mylar films (6 µm, SPEX, Fisher) within a metallic support. For ligand addition experiments, only the bottom (small aperture) side is covered with mylar (though the wells are not individually sealed), leaving the top side (large aperture) open (Figure 1). Samples were briefly stored and transported to the beamline in a humidified box. At the beamline, the open side faces the Microdrop ejector for droplet delivery. After completion of data acquisition, the PEI was transferred to the calibration setup to evaluate droplet ejection.

**Figure 1:**
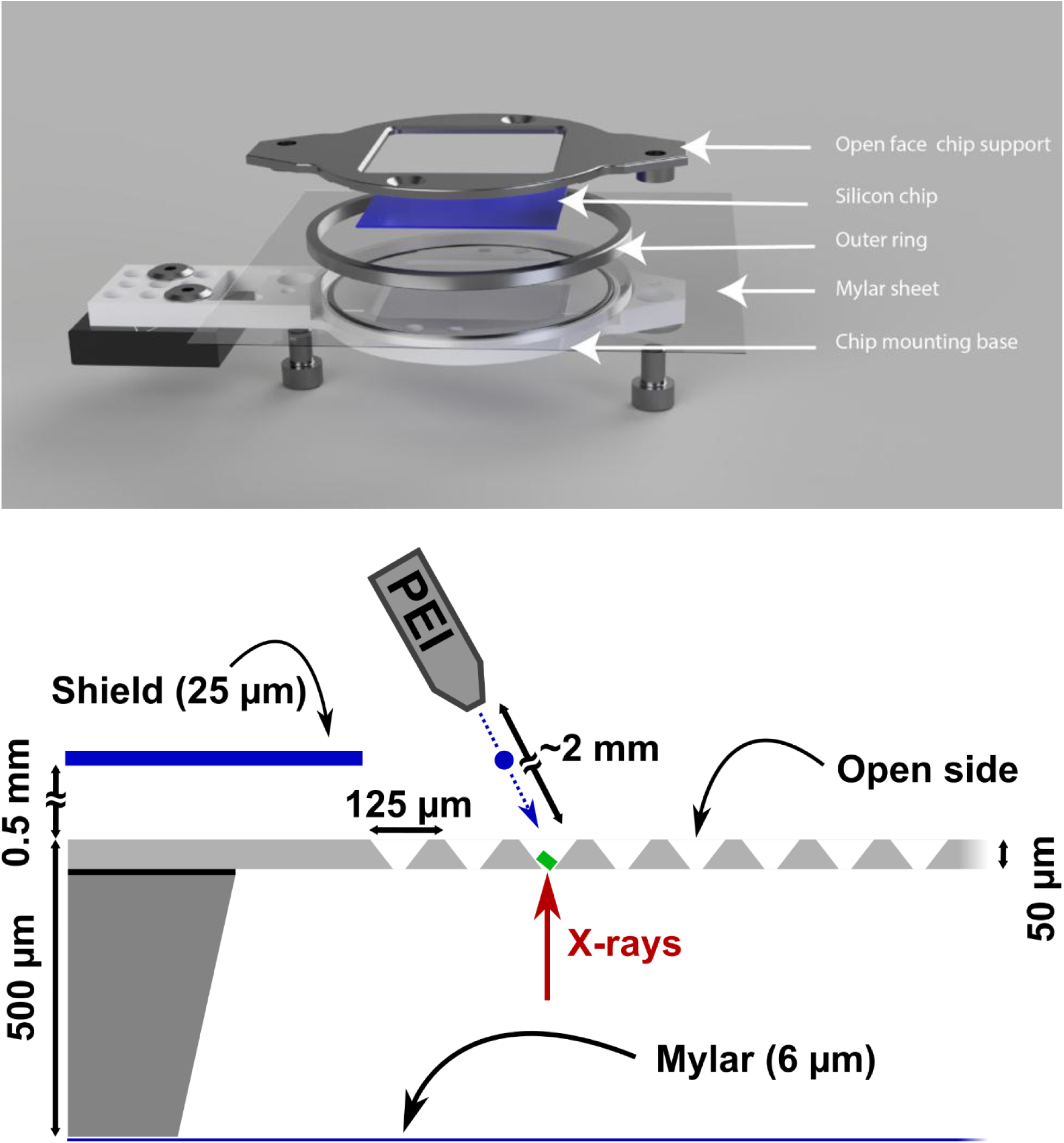
A drop on fixed target sample delivery method for tr-MX experiments. (Top) Exploded view of chip mounting for drop on fixed target experiment. A 6 µm mylar sheet is applied on the chip mounting base and secured with the outer ring. This mitigates sample dehydration from below. The silicon chip sitting on the mylar sheet is then secured with the open face chip support. Once the chip is ready it can be easily mounted on the stages via the kinematic mount. (Bottom) Schematic representation of the side view of the chip, highlighting the open and closed side and the distance between the mylar seal and the wells. The transparent plastic shield is located at about 0.5 mm from the open side chips surface and has a 5 × 5 mm hole to allow droplets to pass through. The figure is drawn to scale.

### 2.2 Drop on fixed target setup

#### 2.2.1 Installation of the setup on the beamline

The drop on fixed target system builds on reported designs (Sherrell *et al*., 2015; Lučić *et al*., 2022; Moreno-Chicano *et al*., 2022, 2019), using high-precision motorised three-axis stages (SmarAct), controlled using a DeltaTau Geobrick LV-IMS-II, an on-axis viewing (OAV) system with a motorised backlight, and a metallic holder for silicon chips (Figure 1 and Figure 2). Modular components and kinematic mounts allow rapid sample exchange and easy adaptation to different beamlines(Horrell *et al*., 2021). The chip’s open side faces downstream of the X-ray beam, and is covered with a film (∼170 µm) with a ∼500 µm gap between the film and the chip. A fixed 5 × 5 mm^2^ opening at the X-ray interaction region enables droplet addition onto the chip while minimising dehydration of the rest of the chip. Absorbent pads (Figure S2) on the fixed frame and chipholder passively increase local humidity to mitigate sample dehydration. Passive humidification was preferred over active humidity control (*i.e.* using a humidified air stream) as the latter frequently resulted in condensation on the chip surface, adding liquid and likely contributing to cross-well contamination (see Results).

**Figure 2:**
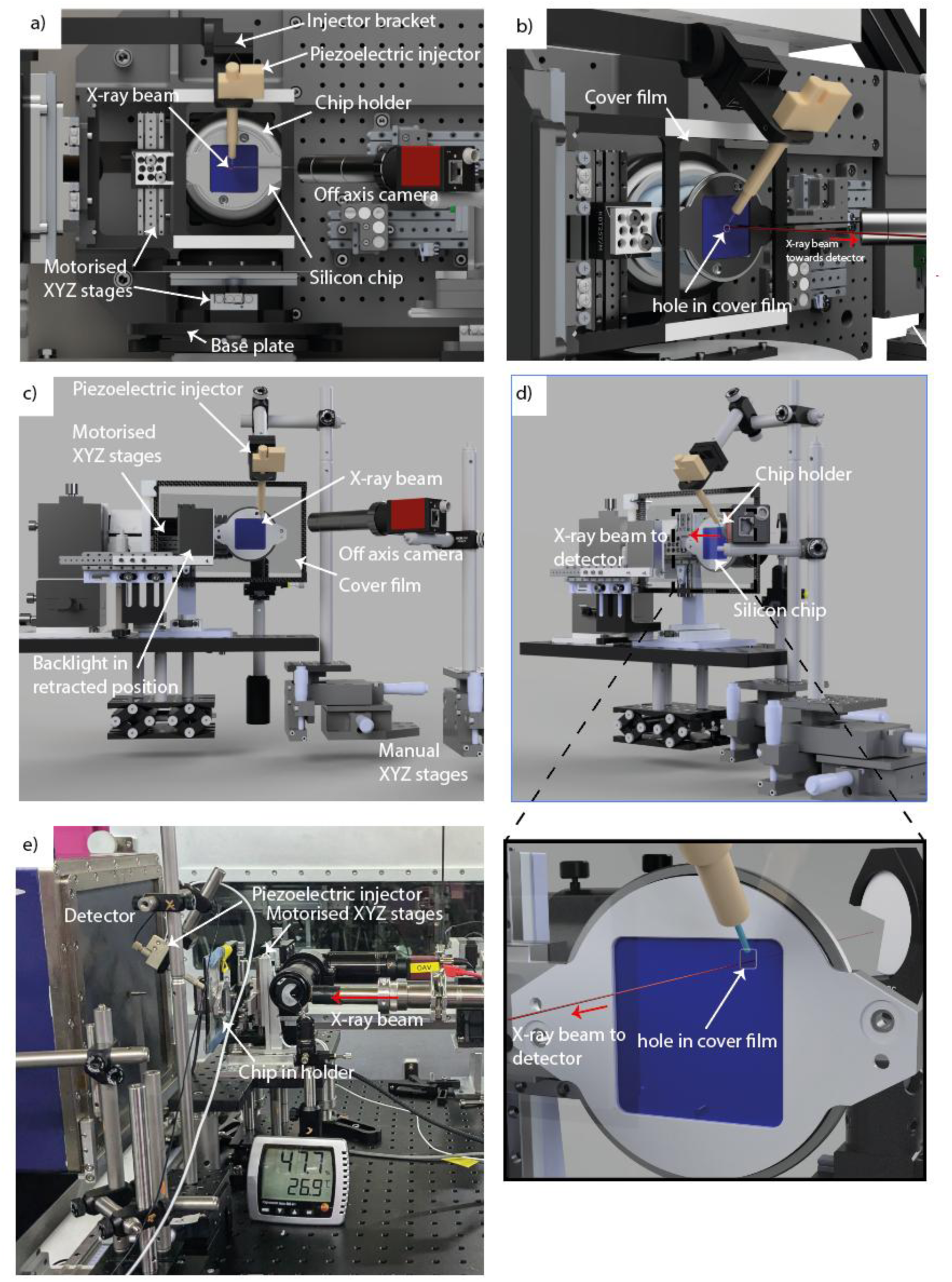
Drop on fixed target setup at various endstations. A typical setup consists of a fixed target chip, containing an open face, which is covered by a cover film that is held in place by a frame. An off axis camera is used to align the droplet ejection with the wells. a) Rendered view of the drop on fixed target setup at I24 Diamond Light Source (DLS) observed from the detector position looking upstream into the X-ray beam; b) rendered view of the drop on fixed target setup at I24 DLS from the side, highlighting the hole in the cover film through which droplets can be added to the fixed target chip; c) rendered view of the drop on fixed target setup at the NCI endstation at PAL-XFEL; d) rendered view from the side of the setup at the NCI endstation at PAL-XFEL; e) photograph of the drop on fixed target setup at the NCI endstation at PAL-XFEL.

Similarly to reported methods (Mehrabi, Schulz, Agthe *et al*., 2019), a piezoelectric injector (PEI), Autodrop Pipette (AD-KH-501-L6; 50 µm inner diameter microdrop Technologies GmbH https://www.microdrop.de/) was used to generate droplets on-demand. By adjusting the pipette’s inner diameter and the triple-pulse waveform parameters, droplets of appropriate size (40–90 pL) are generated (verified by measuring spherical droplet diameter in stroboscopic video). Our testing was done with 10, 30, and 50 Hz dispensing frequencies, though it is compatible with higher frequencies. Dispensing stability and droplet volume were prioritised during optimisation, as evaluated by camera footage; droplet velocity was considered less critical. Importantly, these volumes are smaller than the well volume to avoid overfilling. The PEI mounts via a kinematic bracket to the vertical goniometer table (I24 DLS, Figure 2 a, b) or a Thorlabs post (NCI PAL-XFEL, Figure 2 c, d), and is tilted by 60° relative to the chip. This mounting enables quick switching between experimental and calibration setups; the latter using a stroboscopically operated light and camera to verify and potentially adjust ejection parameters (pulse lengths and amplitudes (*i.e.* voltages); Figure S3). The triple-pulse waveform mode of the dispenser was used, as it provides more control and produces smaller droplets, than the single-pulse mode (Zhang *et al.*, 2024), as validated empirically. The width and amplitude of the second (“expansion”) pulse in the triple-pulse waveform, has the largest impact on droplet volume and velocity. Once optimal parameters are established the PEI is mounted onto the experimental setup at the beamline (Figure 2).

#### 2.2.2. Setup calibration / alignment

The motorised stages align the chip to the X-ray beam position using fiducial markers etched in the chip corners (Sherrell *et al*., 2015), establishing the chip’s coordinate system, correcting any roll, pitch, or yaw relative to the X-ray beam.

Subsequently, the injector is aligned with the X-ray beam using an off-axis camera (Figure 2a and 2c) to visualise droplets landing on the chip. The position of the PEI is adjusted using either the manual three-axis stages at PAL-XFEL or the motorised axis at I24 to ensure that the droplet hits the correct position. PEI alignment is checked before each data collection, as alignment is crucial for droplet ejection accuracy.

### 2.3 Data collection strategy

#### 2.3.1. Collection approach

To enable time-resolved experiments, two dispensing strategies were explored to generate defined time delays (or reaction time; Figure S4 and S5). Critically, droplets are added to every other well in a checkerboard pattern, creating two, fully interleaved datasets (a control and a timepoint) across the entire chip (Figure S6). The interleaved dispensing pattern is programmed into the SmarAct controller.

**“Add & Collect”**: droplets are added to alternating wells, followed by X-ray exposure after a short delay (Figure S4). The attainable time resolution (*i.e.* delay range) depends on the X-ray source. **“Add then Revisit”**: droplets are first added to alternating wells covering 2, 4, 6, 10 or 20 rows within a city block (Figure S5). Subsequently, each row is “revisited” and probed with X-rays. This yields longer, distinct time delays, that can be partially fine-tuned by changing the global acceleration (GA) of the stages. Because the chip is scanned twice, total acquisition time is doubled compared with a ground state (*i.e.* no droplet) experiment.

#### 2.3.2. I24 endstation (DLS)

Room temperature data were collected using an unattenuated 12.8 keV, a 7 × 7 µm^2^ X-ray beam and a Pilatus3 6M detector at 320 mm distance. A flux of 5.4 ⋅ 10^12^ photons s^-1^ was measured at the sample position. As the stages move to each well, a Transistor-Transistor Logic (TTL, 5 V) signal from the Geobrick controller simultaneously triggers the PEI droplet dispensing and detector acquisition (Figure S7). The delay time (or incubation/mixing time) starts when the TTL reaches the PEI and ends when a single frame is recorded (Figure S8). The PEI tip is positioned close to the chip (∼2 mm, Figure 1); with droplet velocities of ∼1 m/s, the time of flight is ∼2 ms. The delay time is not adjusted for droplet time of flight and should be accounted for separately. The resolution of the time delay (for Add & Collect) is set by the detector acquisition time or pulse duration (*e.g.* through the use of a chopper). Due to the quasi-continuous beam available at I24, the detector sets the time delay resolution; *i.e.* 10 ms for the Pilatus3 6M used in this study.

#### 2.3.3. NCI endstation (PAL-XFEL)

Data were collected using an unattenuated 9.5 keV, 3 × 3 µm^2^, 30 Hz, 25 fs (800 µJ / pulse) X-ray beam and a Rayonix MX225 HS (4 × 4 binning) set at 120 mm distance (∼1.77 Å inscribed circle). The SmarAct stages “follow” the X-ray pulses via external TTL signal 1.2 ms after the X-ray pulse (Figure S7B/C). The X-ray pulses are delivered either at 30 Hz (25 fs pulse) or at 10 Hz (25 fs pulse). Here, time delay resolution is set by X-ray pulse duration (25 fs), much shorter than the incubation times. The time between each X-ray pulse, which can be adjusted, defines the maximum incubation time for Add & Collect.

### 2.4. Data processing

The data acquired at I24 (DLS) and NCI (PAL-XFEL) were indexed and integrated using *DIALS* (version 3.23.0) (Winter *et al*., 2018; Beilsten-Edmands *et al*., 2024). Resolution cut-offs were determined based on multiplicity (>10) and monotonic decrease in CC_1/2_. Overall data quality was evaluated by removing an aromatic residue (*i.e.* lysozyme Tyr53; CTX-M-15 Tyr240; and AmpC_EC_ Tyr112) and inspecting the *F*_o_-*F*_c_ map after phasing for reappearance of electron density consistent with the missing residue (see Figure S9, von Stetten *et al*., 2025).

Phases were calculated by molecular replacement with Phaser (McCoy, A. J. *et al*., 2007; PDB 7BHK for lysozyme (Butryn *et al*., 2021), PDB 7BH3 for CTX-M-15 (Butryn *et al*., 2021) and PDB 6T3D for AmpC_EC_ (Lang *et al*., 2020, with ligands removed). Iterative cycles of refinement were performed using Phenix (Adams *et al*., 2002) and manual model rebuilding in Coot (Emsley *et al*., 2010). Avibactam restraints were calculated with eLBOW in Phenix (Moriarty *et al*., 2009).

Evidence for ligand binding was obtained by evaluation of *mF_o_-DF_c_* Polder OMIT maps (Phenix) and *F_o_-F_o_* isomorphous difference maps between time point and resting state structure (Phenix). Subsequently, electron density maps were analysed for any potential cross-well contamination by inspecting *F_o_-F_o_* isomorphous difference maps of the control and the ground state structure.

## 3 Results

### 3.1. Setup design and parameterisation

Drop on fixed target mixing integrates a piezoelectric droplet generator with silicon chip-based approach (Figure 1) (Owen *et al*., 2017; Mueller *et al*., 2015, Mehrabi, Schulz, Agthe, *et al*., 2019). Each chip contains a grid (8 × 8) of “city blocks”, and each block holds a grid (20 × 20) of wells that captures microcrystals. The minimum crystal size retained on the chip depends on the aperture size of the wells (*e.g.* 15, 10 µm). The inter well distance and overall geometry of the wells remains the same for each design (Figure S1). The well volume varies with the chip aperture size (Figure S1), which in turn constrains the volume that can be added to each well (between 122–199 pL; Table S1).

Droplets were added to microcrystal-filled wells on a fixed target chip using a commercial Autodrop Pipette (Microdrop Technologies; 50 µm inner diameter nozzle) as reported (Mehrabi, Schulz, Agthe *et al*., 2019), operated in triple-pulse mode yielding small and stable ejecting droplets (40 – 90 pL) as verified using a stroboscopic camera setup. Dispensing parameters were carefully optimised for each substrate/ligand concentration and dispenser head, prioritising volume and stability. Droplet velocity, accuracy, volume and stability were all affected by the 8 parameters defining the waveform. The second pulse strongly affected velocity and volume (Figure S3).

Dispensing liquid through nozzles with different inner diameters (*i.e.* 30 and 70 µm) was explored. Larger nozzles risk overfilling wells due to the dependence of droplet volume on nozzle diameter, potentially leading to cross-well contamination, while smaller nozzles increases the chance for (irrecoverable) clogging. Initial ejection stability (temporal and spatial) was evaluated over a ∼1 mm distance when optimising parameters. Using optimised parameters (Figure S3) the Autodrop Pipette was operated at the beamline (Figure 2). Alignment began by positioning the chip with respect to the X-ray beam, followed by alignment of the PEI with the first well on the chip.

The silicon chip is mounted in a metal holder connected to a motorised *xyz* stage via kinematic mounts (Sherrell *et al*., 2015). An offline setup with a fast camera (Photron NOVA S12, 12X zoom, Navitar) operated stroboscopically or at high framerates (>7200 Hz), was used to evaluate droplet deposition onto a “dry” chip. Video analysis revealed residual motion and positional overshoot upon arrival at the desired location (Supplementary video 1 and 2), which was dependent on stage stiffness, calibration, and especially acceleration and deceleration parameters (*i.e.* global acceleration (GA)) of the motorised stages (Figure S10). Cumulative residual motion was quantified in the *x* and *y* directions but extended to a lesser extent to the *z*-direction. Optimal GA values balanced well-to-well travel time against positional accuracy, accepting a marginal increase in travel time for higher precision. The GA setting indirectly provides limited tunability of the time points for Add and Collect strategy (see below), by affecting stage-movement delays.

The PEI is positioned downstream of the chip, casting a shadow that varies with detector distance and nozzle angle (Figure 2). Although shallower dispensing angles (*e.g.* 45°) were tested, a more perpendicular angle (*i.e.* 60°) increased the target size and reduces the impact of residual motion in the *z*-direction (Figure S11) with minor detector shadowing. This configuration maintains high (>99.5%) well-hit ratios across the whole chip, as determined through stroboscopic video analysis and by scattering signal from water droplets, dispensed on a dry silicon chip (Figure S12). The incorrectly hit wells (<0.5% of total wells), were clustered at the bottom of the chip, possibly due to larger residual motion in this area.

Optimal loading of the microcrystal slurry onto the silicon chip is crucial for time-resolved experiments, because excess liquid can cause neighbouring well contents to mix via capillary action. This allows reactions to be initiated in multiple wells simultaneously, while X-ray probing occurs one well at a time. A microcrystal slurry is applied to the “top” of the silicon chip (larger, open side of tapered wells; macroscopically smooth side) which is mounted on a sample loading station (Figure S13), in a humidity-controlled enclosure (Horrell *et al*., 2021). A vacuum is applied to the “bottom” of the chip (small aperture side) removing excess liquid, while a sufficiently high relative humidity (>80%) prevents dehydration. The chip is inspected under a microscope; any remaining liquid can be removed by reapplying vacuum or blotting. The semi-open chip design requires careful humidity control at the setup, achieved by localised shielding and installation of wet padding (“passive humidification”). Strong humidifier streams (“active humidification”) are avoided as they can lead to condensation on the chip, which contributes to cross-well contamination (see below).

### 3.2. Generation of time points

Two approaches generate different time points: (i) the “Add and Collect” approach (short time points), where a droplet is added and data acquired before moving to the next well (Figure S4); and (ii) the “Add and Revisit” approach (longer time points), adding (co-)substrate/ligand to multiple wells then revisiting them sequentially for X-ray irradiation (Figure S5). Time points depend on well-to-well travel time, electrical signalling, and droplet time-of-flight and data collection frequency (Figure S8). The setup uses a Geobrick for communication via TTL signals, introducing a stochastic delay (*i.e.* 150 µs inaccuracy). Remounting the chipholder after exchanging a chip alters the Autodrop Ejector to chip distance, changing droplet time-of-flight (∼300 µs inaccuracy). The temporal error, estimated <1 ms for both dispensing approaches, is not account for when setting time delays; it compares favourably to the diffusive properties of small molecules, such as avibactam in CTX-M-15 crystals (4.18 ms; 5.65 ⋅ 10^-6^ cm^2^ s^-1^; 15 × 15 × 5 µm^3^; 50% concentration at centre of crystal; supplementary information; Schmidt, 2020). Thus, the temporal error is negligible compared to the accessible time points.

### 3.3. Interleaved control

To validate fixed target time-resolved experiments a “checkerboard” interleaved control protocol was implemented during data collection (Figure S6). Data from control wells, which received no droplet, covering 50% of the chip, were processed and analysed separately, then compared with ground state structures (*i.e.* no (co-)substrate/ligand added) to reveal potential cross-well contamination. Cross-well contamination in the control wells indicates the main dataset is also contaminated, introducing an undefined incubation time, which inaccurately reflects the anticipated time point. To evaluate the sensitivity of this approach to contamination, synthetic datasets were created, randomly mixing GlcNAc premixed with HEWL data in with apo Hen Egg White Lysozyme (HEWL) data, with varying ratios (*i.e.* 5-75%; Figure S14). Features in the difference maps (*i.e.* appearance of positive density) indicate HEWL•GlcNAc complex formation in part of the data.

Although the control wells provide a limited buffer zone, contamination in these wells implies that more extensive spread cannot be ruled out. Consequently, whenever contamination was observed in the control dataset, all the data from that chip were discarded from the tr-SSX or tr-SFX analysis. Data were collected from city block to city block down and up across the chip in a serpentine manner, suggesting directionality of data collection by itself did not affect cross-well contamination in this setup. Although use of the interleaved control doubles the amount of sample required, its inclusion was critical for the integrity of the time-resolved experiment.

### 3.4. Enzyme samples

To demonstrate the application of this methodology, we studied three enzymes: HEWL and two bacterial serine ß-lactamases (SBLs), CTX-M-15 and AmpC_EC_. HEWL is a model system frequently used for crystallographic method development; it binds different ligands, including GlcNAc (Mehrabi, Schulz, Agthe *et al*., 2019; Butryn *et al*., 2021). CTX-M-15 is a class A extended-spectrum SBL found globally in multiple bacteria, where its production contributes to resistance to many ß-lactam based antibiotics (Castanheira *et al*., 2021). AmpC_EC_ is a class C SBL that is chromosomally encoded by *Escherichia coli* strains. Production of AmpC_EC_ confers resistance to penicillins and cephalosporins (Jacoby, 2009). Co-administration of ß-lactam antibiotics with a ß-lactamase inhibitor, such as avibactam, is a validated method to overcome resistance (Tooke *et al*., 2019). Understanding the mechanism of SBL inhibition by avibactam in a time-resolved manner will lead to improved designs for new antibiotics and/or SBL inhibitors.

#### 3.4.1. Lysozyme

Binding of GlcNAc (221 g/mol, as 226 mM stock in PEI; 76-136 mM final in well) to HEWL was tested at I24 DLS, after mixing for 1.9 s. Final concentration estimates of GlcNAc were based on average droplet size and the average residual volume left per well after chip loading, determined offline using a stroboscopic camera and LED, and estimated by weighing the chips in repeated loading tests. Alongside time point datasets, a ground-state apo-HEWL and a fully equilibrated state were collected; in the latter case GlcNAc was premixed with HEWL for 10 minutes before loading. The data collected on apo-HEWL and the premixed HEWL•GlcNAc complex extended to 1.65 and 1.64 Å resolution, respectively. The time-resolved HEWL•GlcNAc dataset obtained after a 1.9 s delay (Figure 3), using the “Add and Revisit" approach, was refined to 1.69 Å resolution. The 1.9 s mixing dataset (4 revisited rows, GA set to 8) showed strong electron density features in the *mF_o_-DF_c_* polder OMIT map (Figure 3 b) and *F_o_^1.9s^-F_o_ ^apo^* isomorphous difference map (Figure 3 c). The strong features were unambiguously assigned to GlcNAc with an occupancy of 75% (Figure 3 a).

**Figure 3:**
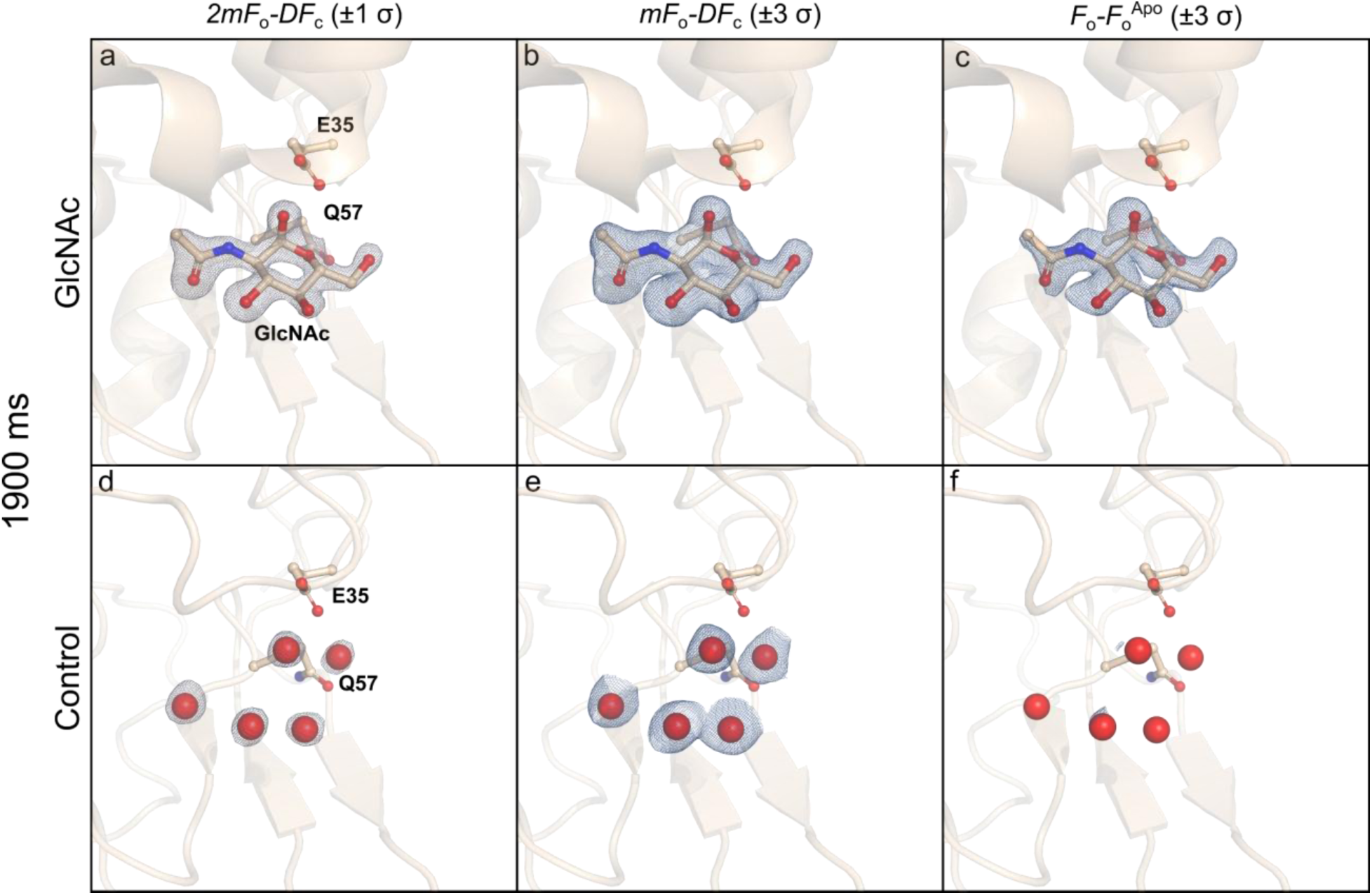
Electron density map analyses of *N*-acetyl-D-glucosamine (GlcNAc) added to Hen Egg White Lysozyme (HEWL) microcrystals obtained through drop on fixed target tr-SSX. Tr-SSX maps for HEWL mixed with GlcNAc (226 mM) for 1.9 s (a - f), collected using the hit-and-return approach at I24 Diamond Light Source determined to 1.69 Å resolution. a) *2mF_o_-DF_c_* maps of the active site of the HEWL•GlcNAc complex (radius: 1.5 Å, contour: 1 σ); b) *mF_o_-DF_c_* polder OMIT map (radius: 1.5 Å, contour: 3 σ; integrated density: 63.1 e^-^ Å^-3^); c) *F ^1.9s^-F ^Apo^* isomorphous difference map (contour: 3 σ); d) *2mF_o_-DF_c_* map of the interleaved control (*i.e.* without ligand added; radius: 1.5 Å, contour: 1 σ); e) *mF_o_-DF_c_* polder OMIT map of the interleaved control (radius: 2.0 Å, contour: 3 σ); f) *F_o_^Control^-F_o_^Apo^* isomorphous difference map (contour: 3 σ; integrated density: 0.67 e^-^ Å^-3^).

Careful analyses of the *mF_O_-DF_C_* polder OMIT electron density map (Figure 3 e) from interleaved control data, revealed water molecules at the GlcNAc binding site, with no evidence of active site bound GlcNAc. Additionally, *F_o_^Control^-F_o_ ^Apo^* isomorphous difference maps (Figure 3 f) agree with the *mF_o_-DF_c_* polder OMIT maps, confirming the absence of GlcNAc ligand binding.

The GlcNAc premixed data indicates full occupancy of GlcNAc in the HEWL structure (Figure S15). Comparison of an isomorphous difference map calculated between the GlcNAc premixed and 1.9 s mixing time point data sets (Figure S15 d), manifests additional electron density, consistent with incomplete occupancy after 1.9 s mixing.

#### 3.4.2. CTX-M-15

Using the approach at I24 DLS and at NCI PAL-XFEL, various time-resolved datasets after avibactam addition (265 g/mol, 200 mM stock; 66-100 mM final) to CTX-M-15 were collected. Avibactam binds to and covalently-modifies the active site Ser70 within 80 ms (partially; PAL-XFEL; Add and Collect) and 2.6 s (fully; I24; Add and Revisit (4 revisited rows; GA = 10)) incubation times (Figure 4). Diffraction data at 2.6 s incubation time extended to 1.65 Å, and provided clear evidence for the presence of two avibactam molecules: one at, and one close to, the active site, as indicated by pronounced positive features in the *mF_o_-DF_c_* polder OMIT map. Refinement suggests full occupancy for an avibactam ligand in its ring-opened state covalently bound to Ser70; and a second, intact, non-covalently bound ligand near the active site that refined to 69% occupancy (Figure 4 a-f). The presence of a second avibactam molecule likely reflects the relatively high concentration used (200 mM). The “second” bound avibactam was also present in premixed complex structures, as determined by both traditional, rotation-based cryogenic and room temperature crystallography using fixed targets (Hinchliffe *et al*., 2025). Comparison of the CTX-M-15•Avi complex after 2.6 s incubation time with data for uncomplexed CTX-M-15, using an isomorphous difference map (Figure 4 f), provides further unambiguous evidence for the presence of avibactam.

**Figure 4:**
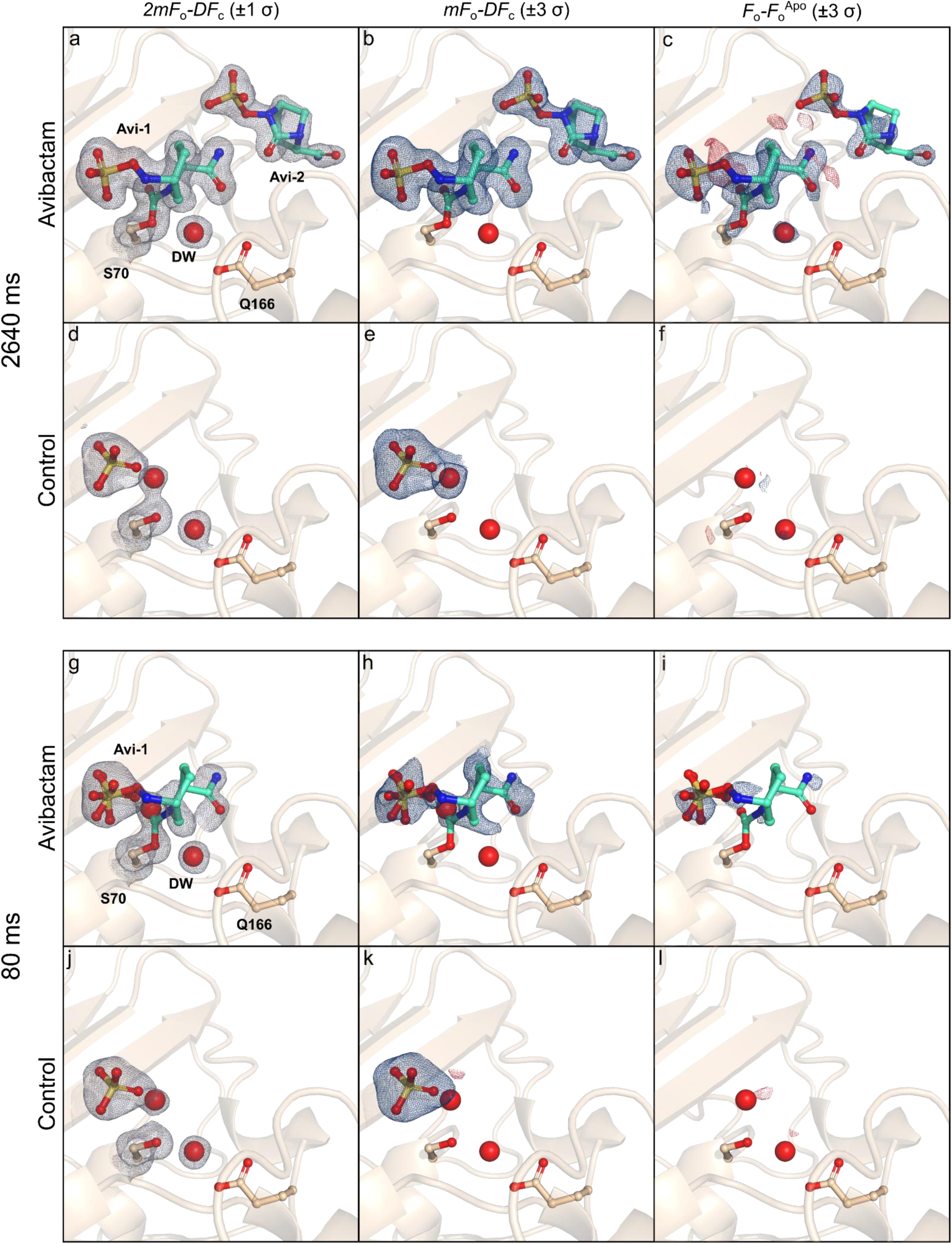
Electron density map analyses of avibactam (Avi) bound to CTX-M-15 microcrystals obtained through drop on fixed target tr-SSX and tr-SFX. Tr-SSX maps for CTX-M-15 mixed with Avi (200 mM) for 2.6 s (a - f), collected using hit-and-return approach at I24 Diamond Light Source, are determined to 1.55 Å resolution. a) *2mF_o_-DF_c_* map of the active site of the CTX-M-15•Avi complex (radius: 1.5 Å, contour: 1 σ); b) *mF_o_-DF_c_* polder OMIT map (radius: 2.0 Å, contour: 3 σ); c) *F_o_^2.6s^-F ^Apo^* isomorphous difference map (contour: 3 σ; integrated density: 40.64 e^-^ Å^-3^); d) *2mF_o_-DF_c_* map of the interleaved control (*i.e.* without ligand added; radius: 1.5 Å, contour: 1 σ); e) *mF_o_-DF_c_* polder OMIT map (radius: 2.0 Å, contour: 3 σ); f) *F_o_^Control^-F_o_^Apo^* isomorphous difference map (contour: 3 σ; integrated density: 1.71 e^-^ Å^-3^). Tr-SFX maps for CTX-M-15 mixed with Avi for 80 ms (g - l), collected using the Eject and Collect approach at NCI PAL-XFEL are determined to 1.50 Å resolution. g) *2mF_o_-DF_c_* map of the active site of the CTX-M-15•Avi complex (radius: 1.5 Å, contour: 1 σ); h) *mF_o_-DF_c_* polder OMIT map (radius: 2.0 Å, contour: 3 σ); i) *F_o_^80ms^-F_o_^Apo^* isomorphous difference map (contour: 3 σ; 6.40 e^-^ Å^-3^); j) *2mF_o_-DF_c_* map of the interleaved control (*i.e.* without ligand added) (radius: 1.5 Å, contour: 1 σ); k) *mF_o_-DF_c_* polder OMIT map of the interleaved control (radius: 2.0 Å, contour: 3 σ); l) *F_o_^Control^-F_o_^Apo^* isomorphous difference map (contour: 3 σ; integrated density 1.43 e^-^ Å^-3^).

The time-resolved data were acquired using the interleaved, checkerboard control strategy (*i.e.* without ligand addition). Chip-by-chip analysis indicated the absence of cross-well contamination, validating each time point, and allowing 3 chips to be merged for the 2.6 s dataset, while the complete 80 ms dataset is from a single chip. The *mF_o_-DF_c_* polder OMIT electron density maps from the control data, showed the potentially deacetylating water (DW) molecule and a sulfate ion at the active site, with no features consistent with the presence of avibactam. The refined active site sulfate ion located at the active site is likely derived from the crystallisation condition, which contains 2 M (NH_4_)_2_SO_4_. The *F_o_^Control^-F_o_^Apo^* isomorphous difference maps are featureless and in agreement with the *mF_o_-DF_c_* polder OMIT electron density maps, unambiguously demonstrating the absence of avibactam.

A fourth chip for the 2.6 s time-resolved datasets for the CTX-M-15•Avi complex was excluded due to cross-well contamination (Figure S16), despite all the chips being prepared in a seemingly identical manner. Analysis of the *mF_O_-DF_C_* polder OMIT electron density maps from the fourth chip and its corresponding control, revealed clear evidence for the presence of ligand in both the control and ligand added datasets, confirmed by *F_o_^Control^-F_o_ ^Apo^* isomorphous difference maps. As anticipated, the *F_o_^Avibactam^-F_o_^Control^* isomorphous difference map (Figure S16 g) displayed no significant difference densities, indicating near-homogenous spread of ligand across the entire chip. No signs of cross-well contamination were apparent prior to analysis of the control data, highlighting the importance of careful analysis of the diffraction data from each chip.

The 80 ms dataset (Figure 4 g-l) demonstrates clear density corresponding to ring-opened Avibactam at the active site. The *mF_O_-DF_C_* polder OMIT electron density map shows the ring-opened Avibactam covalently bound to the catalytic Ser70, albeit at lower occupancy (33%). Notably, no density for a second Avibactam molecule near the active site was observed, consistent with occupancy at the second site after reaction at the active site, providing proof of principle for utility of the method to provide time-resolved information, as explored in detail in the accompanying paper (Hinchliffe *et al*., 2025).

#### 3.4.3. AmpC_EC_

We performed time-resolved experiments on AmpC_EC_ with avibactam (200 mM stock; 66-100 mM final) extending to 3.5 Å resolution (Figure S17), collected at NCI PAL-XFEL. One of the two AmpC_EC_ chains in the asymmetric unit provided evidence for partial occupancy for avibactam bound to the catalytic Ser64. Strong, positive difference density features in the *mF_o_-DF_c_* polder OMIT maps and *F_o_^80ms^-F_o_^apo^* isomorphous difference maps confirm the presence of avibactam. Refinement of the atomic model against the diffraction data suggested an ∼80% occupancy of ring-opened avibactam that is covalently bound to Ser64. In agreement with previous observations (Lang *et al*., 2021), binding of avibactam to Ser64 did not result in substantial changes to the overall fold of AmpC_EC_, nor to the conformation of active site residues. Importantly, inspection of the *mF_o_-DF_c_* polder OMIT map and the *F_o_^Control^-F_o_^Apo^* isomorphous difference map of the interleaved control, suggested that no cross-well contamination occurred during data collection, validating the time-resolved data.

## 4. Discussion

Our setup provides a drop on fixed target sample delivery method capable of performing tr-MX experiments at both synchrotron and XFEL facilities, with, importantly, an interleaved control strategy applied to every chip. The established fixed target approach deploying silicon chips at I24 DLS is expanded with the capacity for liquid mixing. The achievable minimal mixing times depend on the light source, *i.e.* pulsed XFEL or continuous synchrotron, and the data collection approach, *i.e.* ‘Add and Collect’ or ‘Add and Revisit’ strategies. These approaches provide opportunities for time-resolved datasets acquired at time points between 2-80 ms, and distinct timepoints (*i.e.* 1.9, and 2.6 s; depending on the GA), with high temporal accuracy. Direct comparisons of time-resolved data using the drop on fixed target approach with other methodologies is difficult because experimental conditions vary widely. Nevertheless, comparison suggests that different sample delivery methods are likely to produce different results. For instance, a comparison of the time-resolved HEWL data presented here (specifically the 1.9 s time point), and those reported using a drop on drop on demand approach (2 s time point, (Butryn *et al*., 2021)), suggests a slower progression of GlcNAc binding to HEWL in the drop on fixed target setup based on ligand occupancy analysis, despite using a higher final ligand concentration. Although the underlying mechanism for mixing dynamics has not been explicitly characterised in our work, several possible explanations can be proposed. The drop on drop on demand approach relies on repeated deposition of small droplets onto a larger droplet which also induces turbulent mixing, enhancing the rate of diffusion. In contrast, the drop on fixed target method involves deposition of a single droplet onto a partially filled well, where some of the energy of the impact is partially dissipated by the walls of the solid support, which could reduce turbulence and slow down the mixing process. Alternatively, removal of liquid from each aperture could cause partial crystal dehydration, potentially leaving a viscous layer of solutes (*e.g.* [PEG] or salt), impacting mixing efficiency. Although the exact mechanism remains to be identified, these observations underscore the importance of exercising caution when comparing time points across different sample delivery strategies for diffusion-dependent reactions. Instead, current best practices for evaluating tr-MX results are best restricted to comparing internally consistent time series (*i.e.* using the same sample delivery system throughout) to ensure reliable and meaningful interpretation of the particular reaction coordinates.

The interleaved control (*i.e.* without ligand added) was necessary to evaluate potential cross-well contamination, which could lead to premature initiation of a reaction, invalidating the putative time-resolved series. Cross-well contamination could occur due to droplet ejection inaccuracy, droplet volume overfilling the well, or the presence of excess mother liquor (or water from humidification) on the chip after loading. Although droplet volume (40-90 pL) and accuracy (>99.5%) are not yet continuously monitored during data collection, the offline setup and dry chip tests provide consistent results, suggesting that droplet volume and accuracy are unlikely sources of contamination in our setup. Stable ejection is confirmed after data collection, consistent with sustained ejection stability throughout the process. The most likely cause of cross-well contamination starts with loading each chip. Failure to remove all of mother liquor from the boundary layers surrounding each well within a city block is more difficult to control and requires careful balancing of suction power against drying out of the crystals. The viscosity of the mother liquor (affected by *e.g.* [PEG]) can impact profoundly on this step, as well as use of chips with small aperture size (*i.e.* 5 µm) compared to larger aperture wells. Fortunately, a checkerboard dispensing pattern provides unambiguous validation of a well-executed time-resolved experiment, and efficiently reveals cross-well contamination. It is important to note that with the current design, even experienced users will encounter cross-well contamination on occasion. Therefore, interleaved controls are akin to dark and light illumination interleaving, as done frequently for laser triggered, time-resolved experiments (Doak *et al*., 2024; Kubo *et al*., 2017; Gotthard *et al*., 2024). Accordingly, interleaving control data with experimental time-point data is considered best practice to validate time-resolved experiments.

Our results focus on droplet-based ligand addition strategies that are very efficient with samples, easy to conceptualize, somewhat more challenging to execute, and absolutely generalizable across most of enzymology. Given the observations concerning cross-well contamination presented here and elsewhere (Gotthard *et al*., 2024) it is important to discuss the implications for other aperture-aligned, chip-based, time-resolved crystallography experiments. For instance, use of photocaged chemicals to create (super) saturated solutions upon illumination is also vulnerable to cross-well contamination by diffusion to neighbouring wells that are inadvertently connected by solution bridges if/when excess liquid is present. Moreover, release of gaseous compounds such as O_2_ or NO from photocaged chemicals could also lead to gas exchange through an interconnected atmospheric pathway between the two Mylar sheets that enclose the chip, but do not hermetically seal each individual well. Although it is difficult to quantify the impacts of these potential sources of contamination, the present study shows reason for caution when deploying fixed-target based methods, and emphasises the need for the implementation of unambiguous, rigorous controls to validate experimental results.

## 5. Conclusion

We have demonstrated the successful design and implementation of drop on fixed target sample delivery method for tr-MX experiments at both synchrotron and XFEL facilities. The effectiveness of the method for time resolved crystallography studies was validated through the successful demonstration of avibactam binding to CTX-M-15 and AmpC_EC_, and of GlcNAc to HEWL. This versatile and robust method utilises a commercially available piezoelectric droplet dispensing pipette to deliver picolitre volumes droplets (40 - 90 pL) onto protein microcrystals that have been immobilized on a solid support, providing a range of reaction time points (10s ms to several seconds), with high temporal accuracy (estimated <1 ms error). This strategy expands the capabilities of fixed target platforms for tr-MX.

Our results highlight the essential role of interleaved controls in validating time points and ensuring integrity of time-resolved datasets. The method directly addresses concerns for cross-well contamination, an issue that may arise inadvertently even for experienced users. We find that potential contamination is likely linked to the challenge of uniformly removing excess mother liquor from the chip during the loading process. Stringent application of these checkerboard controls, akin to established best practices in laser-triggered experiments, provides a framework for validating time-resolved experiments in other aperture-aligned and fixed-target chip-based serial crystallography studies. While interleaved data acquisition doubles the required sample volume, the overall sample efficiency remains favourable when compared to alternative mixing strategies. It offers an attractive alternative to flow-based mixing approaches, particularly for fragile or mechanically sensitive crystal samples. Moreover, it avoids issues commonly associated with jet stability, although droplet flight trajectory is critical to our approach. The successful deployment at both synchrotron (I24, DLS) and XFEL (PAL-XFEL) facilities validates the broad adaptability of this setup.

Future developments will focus on real-time monitoring of droplet ejection to further enhance reliability. This could involve the use of stroboscopic imaging or analysis of background solvent rings to provide immediate feedback during data collection on potential technical issues.

## Acknowledgements

We thank the staff supporting our beamtimes (2022-2nd-NCI-008, 2023-2nd-NCI-014) at NCI, PAL-XFEL. The authors also thank the Global Science experimental Data Hub Center (GSDC) at the Korea Institute of Science and Technology Information (KISTI) for providing computing resources and technical support. We also thank the staff for experiments (MX-25260) at beamline I24, DLS. AMO, RLO, PA, JG, and JJAGK received support from DLS and UK Science and Technology Facilities Council (STFC). AMO is the recipient of a Royal Society Wolfson Fellowship RSWF\R2\182017 and a Wellcome Investigator Award 210734/Z/18/Z. CLT thanks the University of Bath Prize Fellowship Scheme and the UK Medical Research Council for fellowship funding (UKRI330). Research was supported by the BBSRC-funded South West Biosciences Doctoral Training Partnership (training grant reference BB/T008741/1, studentships to LP and MB), and the Interdisciplinary Bioscience Doctoral Training Partnership (grant number BB/T008784/1, studentship to EIF). PAL thanks the National PhD Training Program in Antimicrobial Resistance Research by the Medical Research Foundation (MRF-145-0004-TPG-AVISO) for funding a postgraduate studentship. CLT and JS thank the UK Medical Research Council for support through the grant MR/T016035/1. This work is part of a project that has received funding from the European Research Council under the European Horizon 2020 research and innovation program (PREDACTED Advanced Grant Agreement no. 101021207) to JS. P.R. thanks the Wellcome Trust (227298/Z/23/Z). EIF and CJS were supported by the Ineos Oxford Institute for Antimicrobial Research.

