## Supplementary Information for "Drop on fixed target reaction initiation approach for serial and time resolved crystallography"

[a]Diamond Light Source, Harwell Science & Innovation Campus, Didcot, OX11 0DE, United Kingdom.

[b]Research Complex at Harwell, Rutherford Appleton Laboratory, Didcot, OX11 0FA, United Kingdom.

[c]Current address: European Synchrotron Radiation Facility, 71 Avenue des Martyrs, 38000, Grenoble, France.

[d]School of Biochemistry and Cellular and Molecular Medicine, University of Bristol, University Walk, Bristol, United Kingdom.

[e]Current address: Department of Chemistry & Molecular Biology, University of Gothenburg, Medicinaregatan 7 B, 41390 Göteborg, Sweden.

[f]Department of Chemistry and the Ineos Oxford Institute for Antimicrobial Research, University of Oxford, 3 Mansfield road, Oxford, OX1 3TA, United Kingdom.

[g]Department of Life Sciences, University of Bath, Claverton Down, Bath, BA2 7AX, United Kingdom.

[h]Centre for Computational Chemistry, School of Chemistry, University of Bristol, Cantock's Close, Bristol, BS8 1TS, United Kingdom.

[i]Pohang Accelerator Laboratory, Pohang University of Science and Technology, Pohang 37673, Republic of Korea.

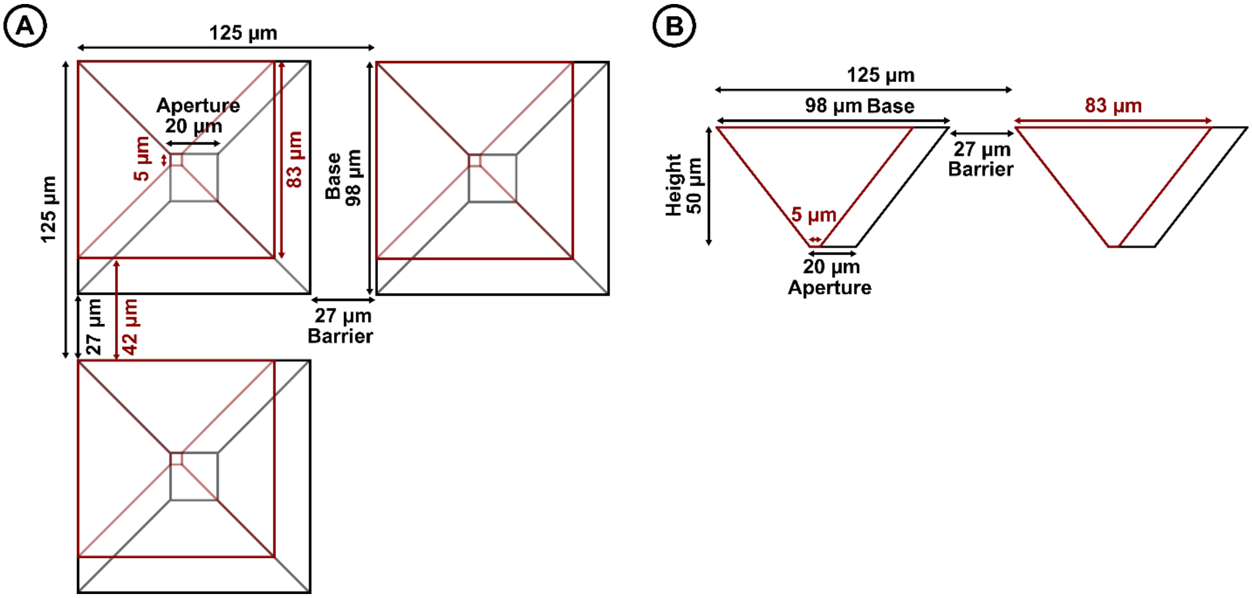

Figure S1: General parameters for the overall design of the Oxford-style silicon chip (ECS Partners Ltd, Southampton, UK). The outlines for a 20 µm and 5 µm aperture size chip are shown in black and red, respectively.

| **Aperture size (µm)** | **Base size (µm)** | **Height (µm)** | **Inter well distance (µm)** | **Barrier length** | **Volume (pL)** |
|---|---|---|---|---|---|
| 20 | 98 | 50 | 125 | 27 | 199.4 |
| 15 | 93 | 50 | 125 | 32 | 171.2 |
| 10 | 88 | 50 | 125 | 37 | 145.4 |
| 7 | 85 | 50 | 125 | 40 | 131.2 |
| 5 | 83 | 50 | 125 | 42 | 122.2 |

Table S1: aperture size, base size, height volume, inter well distance. Volumes were calculated using:

$$V_{well} = \frac{h}{3}(a^2 + b^2 + ab)$$

where *h* is height, *a* is aperture, and *b* is base, all in microns.

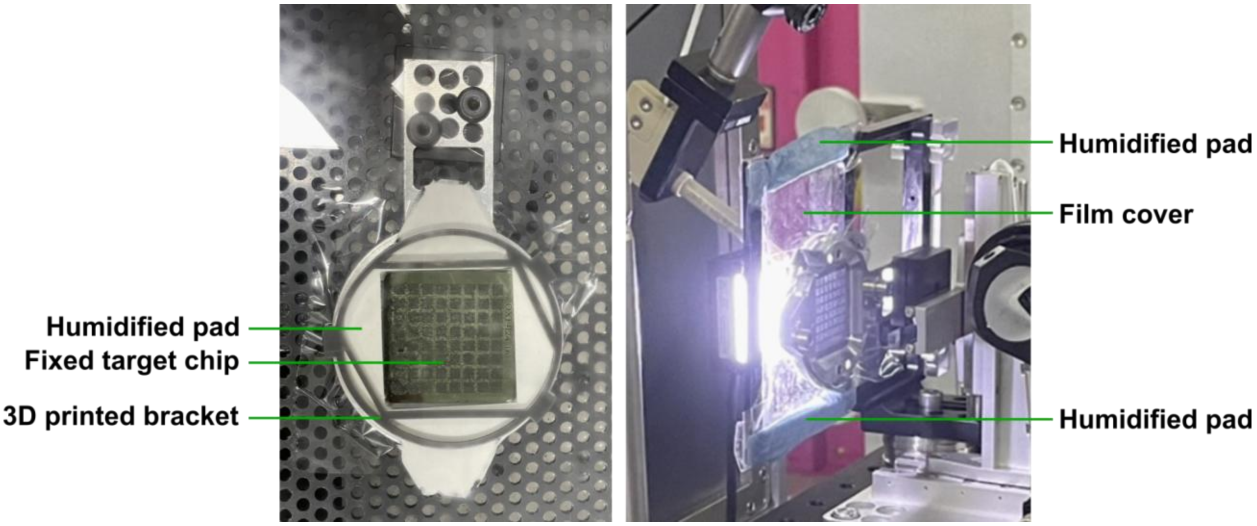

**Figure S2: Passive humidity control during the drop on fixed target experiment.** Left: Fixed target chip, loaded with a microcrystal slurry and mounted in a chip holder, assembled under controlled humidity conditions. The backside of the chip is sealed with Mylar (6 µm), while the open face is surrounded by wetted blotting paper (Thermo Fisher Scientific), held in place by a custom 3D printed bracket. Right: Picture of the setup at the NCI instrument (PAL-XFEL). The open face of the chip is covered with a transparent film fixed in a frame. A small hole ($5 \times 5$ mm$^2$) in the film, allows for droplets to be added onto the wells. Two additional absorbent pads positioned above and below the chip passively increase the local humidity during data collection.

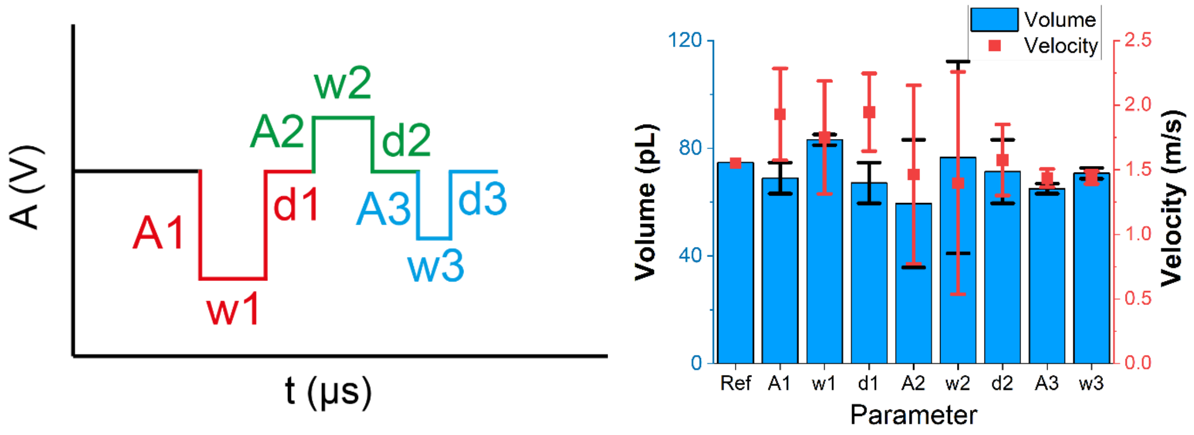

**Figure S3: Parameters constituting the triple pulse actuation waveform used to generate droplet and their impact on the droplet volume and velocity.** Image on the left is a schematic representation of a pulse sequence, not to scale. Reference bar (Ref) refers to manufacturer suggested parameters (*i.e.* A1 = -75 V, w1 = 19 µs, d1 = 13 µs; A2 = 48 V, w2 = 17 µs, d2 = 6 µs, A3 = -32 V, w3 = 20 µs), which can be obtained from the supplier on demand, for stable ejection of water droplets. Note these are different for individual nozzles. Each of the 9 parameters was individually varied from an upper to lower value between which ejection was stable, starting from the manufacturer suggested optimal parameters for water dispensing. Average volume and velocity are derived from video analysis. These averages are plotted, while the black (volume) and red (velocity) bars represent deviation from the average, illustrating the sensitivity of either volume or velocity to the particular parameter. The different pulses of the actuation waveform are highlighted as follows: red, first "initiation" pulse; green, second "expansion" pulse; third, blue "termination" pulse. Droplet volume was determined from stroboscopically operated camera setup (*i.e.* fixed frame rate for camera, with a stroboscopically operated LED light), where a pixel to µm conversion was used to determine the radius of a spherical drop to calculate the volume. Velocity was determined by determining the distance (using pixel to µm conversion) from the meniscus to the centre of the droplet after ejection over time determined by changing the stroboscopic delay. Note that d3 (left) is the delay in pulse 3 of the waveform that can be set to a numeric value, however as it is not immediately followed by a pulse, it has no effect on the droplet size or velocity, and was therefore not considered for further optimisation.

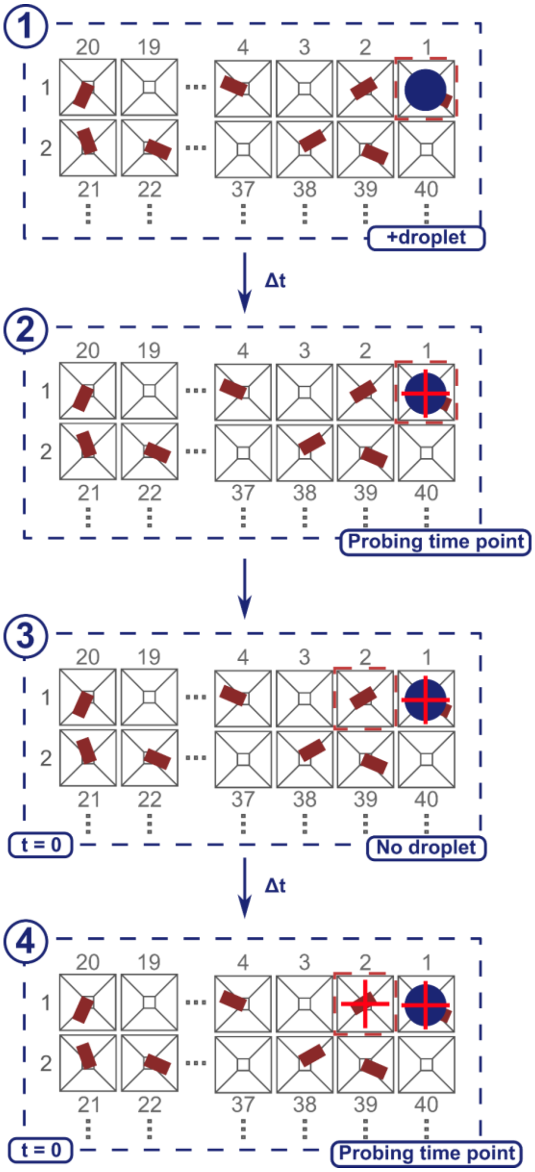

**Figure S4: “Add and Collect” data collection strategy for short time delay tr-SSX and tr-SFX with drop on fixed targets .** The “Add and collect” strategy adds a droplet (blue filled circle) to the microcrystal slurry on the fixed target chip at the well aligned to the X-ray beam (red dashed square) (1). A short delay (Δt between (1) and (2)) precedes data acquisition (red cross) (2). No droplet is added to the neighbouring well when it is aligned to the beam (3); however, the well & sample experiences the same time delay before data acquisition (4).

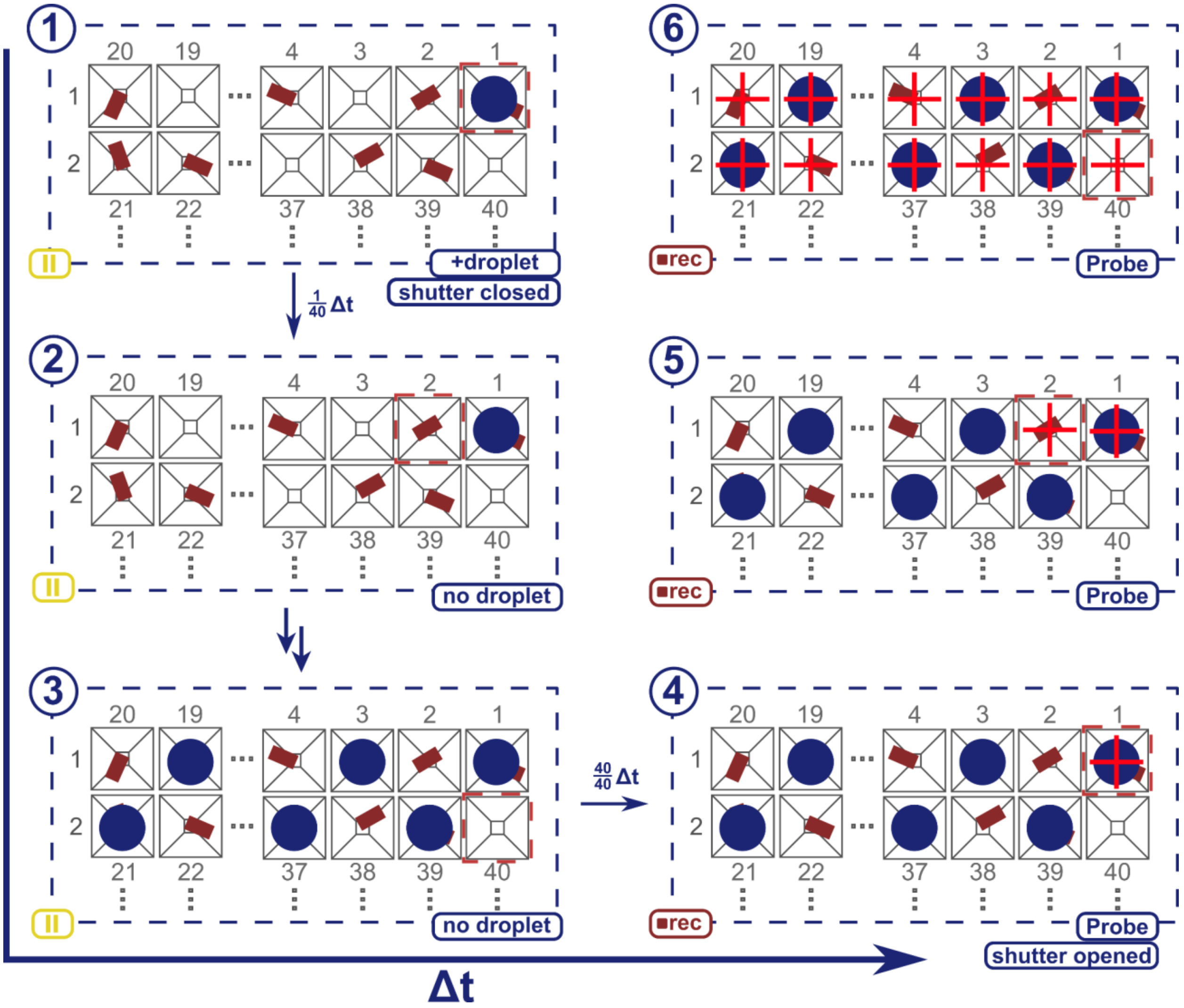

**Figure S5: "Add and Revisit" data collection strategy for drop on fixed target.** The "Add an revisit" data collection strategy, where after closing the shutter, a droplet is added sequentially to every other well for 2, 4, 6, 8, 10, or 20 rows (1, 2, and 3), before opening the shutter (4). Next each well is revisited with X-rays, enabling data collection at set time points (5, 6). Exact time delays can be modulated by changing the number of revisited rows or altering the global acceleration (GA) parameter. The colour scheme is the same as for Figure S4.

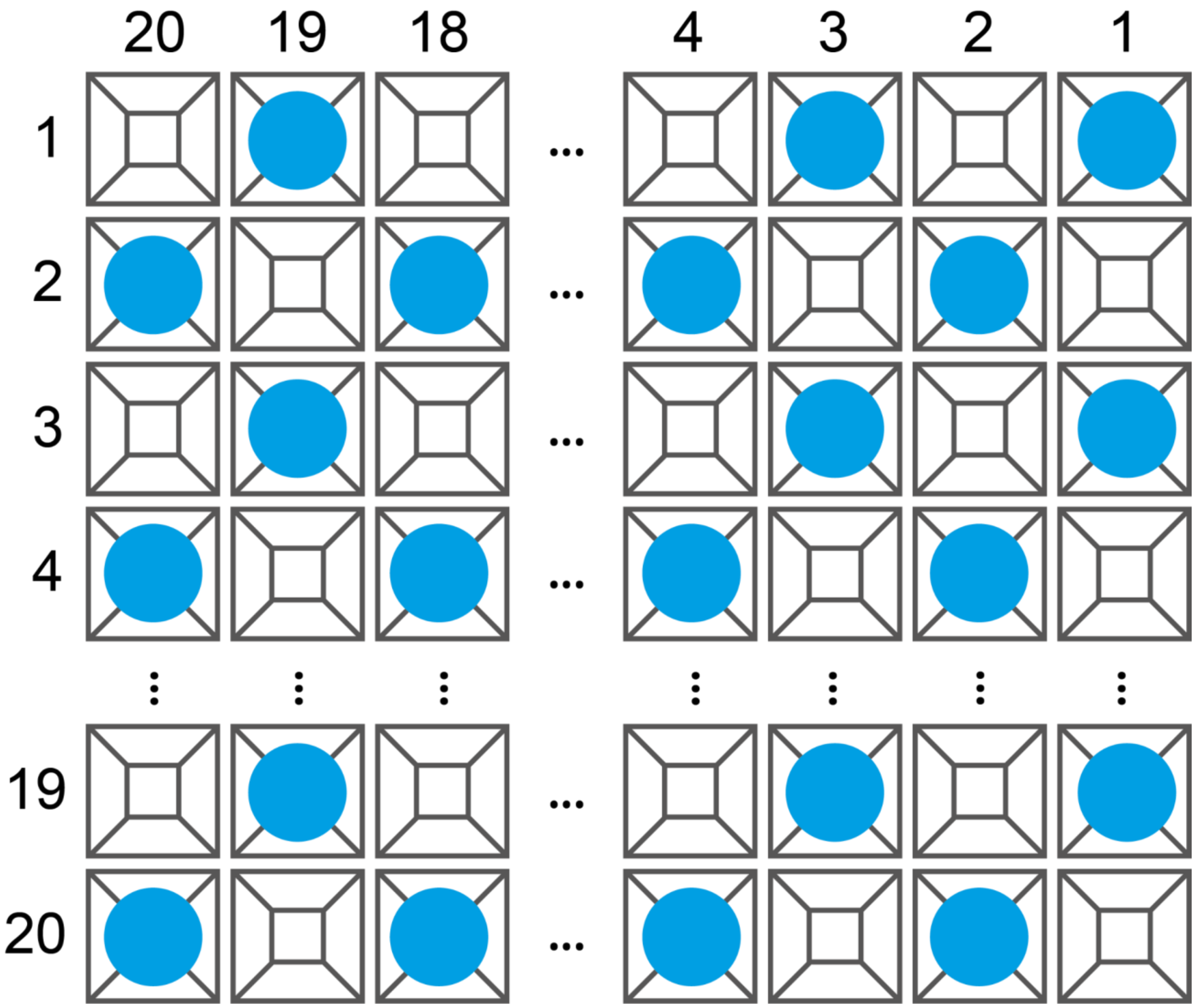

**Figure S6: Checkerboard dispensing pattern to establish an interleaved control during data collection.** Droplets are added to every other well and incubated for the intended time point, while the control wells are left without droplet addition for the same amount of time, prior to X-ray exposure. The interleaved, internal control should always yield a ground state structure.

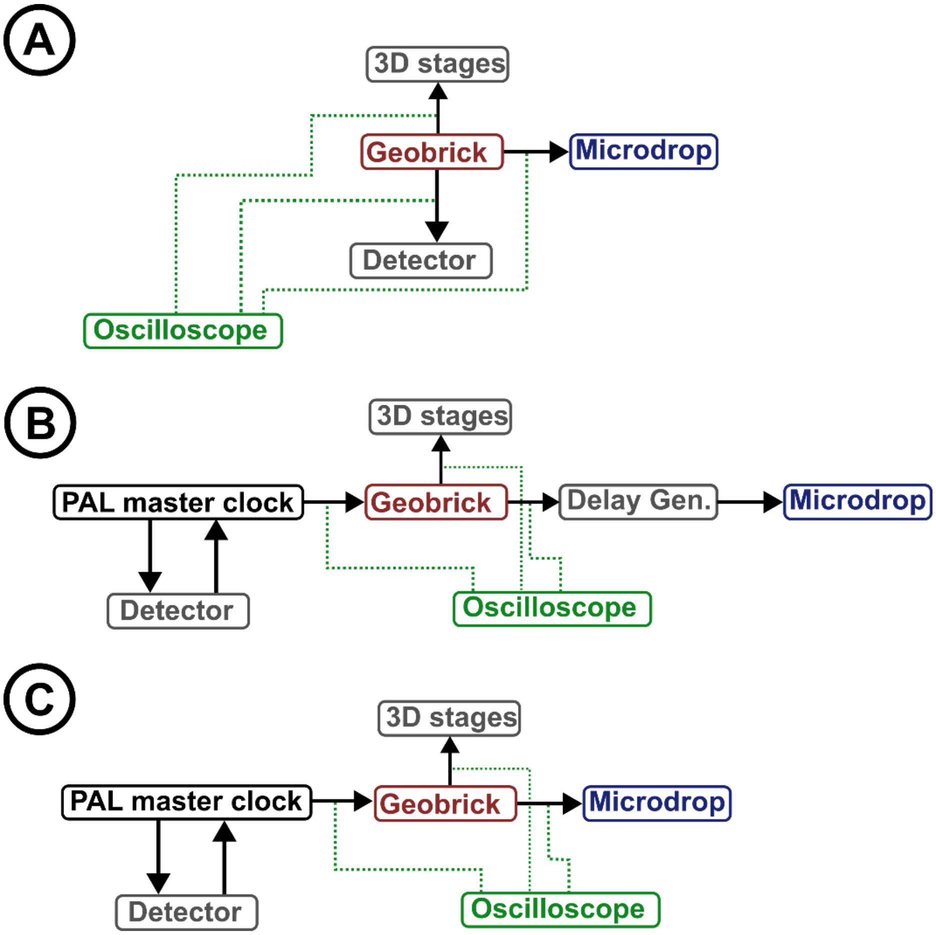

**Figure S7: Schematic signalling diagram of the setup at I24 Diamond Light Source and NCI at PAL-XFEL.** The arrows indicate signaling communication indicating dependency. The green dashed lines indicated monitoring of signals. A) At the start of a run, a transistor-transistor logic (TTL) signal is generated by the Geobrick. This in turn, signals the motorised 3D stages to start movement, and readies the detector for data acquisition, while also providing a signal to the microdrop ejector to eject at the right time. Acquisition of an image is confirmed by the detector, after which the Geobrick restarts the process of signaling the stages to move to the next well. B) "Add and Collect" data acquisition approach, where a droplet is ejected and data is acquired after a fixed delay, set by the delay generator (Delay Gen.). C) "Add and Revisit" data acquisition approach, where the pulse picker is closed, small molecule containing droplets are added to the chip, before opening the pulse picker again and revisiting the wells for data acquisition. Note that programmable code on the Geobrick alters between signalling droplet ejection in one well and no ejection in the next. And note the difference in signalling between I24 at Diamond Light Source and NCI at PAL-XFEL.

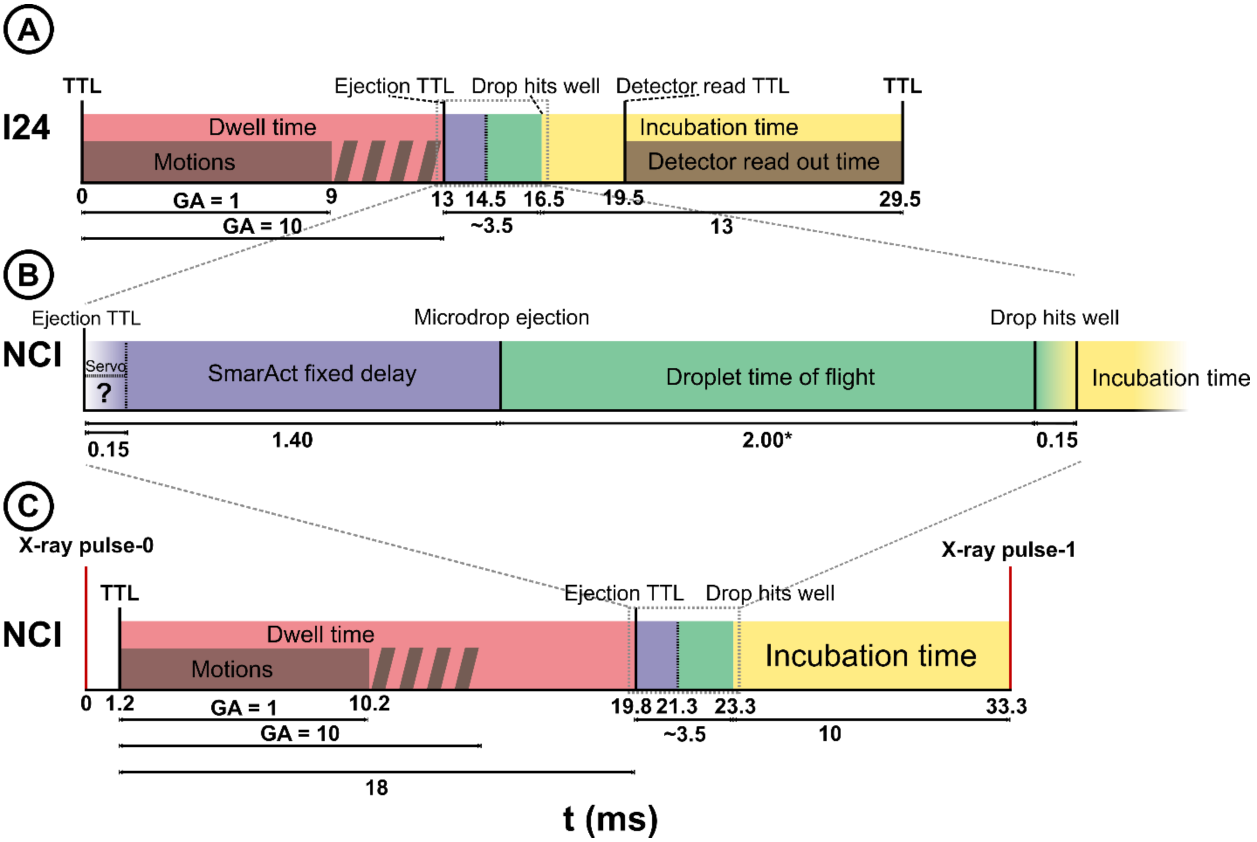

**Figure S8: Signalling and timing overview for "Add and collect" data collection strategy at Diamond Light Source I24 and NCI at PAL-XFEL.** A) At the start of a run, a transistor-transistor logic (TTL) signal generated by the Geobrick to start motions of the motorised stages (GA is the programmable global acceleration parameter for the motors), followed by a TTL signal to the droplet ejector for droplet generation. After a delay, another TTL signal is sent to the detector to start data acquisition. Once completed, the detector communicates with the Geobrick, which restarts the process of signaling the motorised stages to move to the next well. In this example an arbitrary 16.5 ms time delay (*i.e.* incubation time or time point) was chosen. The motions can take up 9–13 ms, depending on what GA is chosen. Note that programmable code on the Geobrick alters between signalling droplet ejection in one well and no ejection in the next. B) Enhanced view of timings from the moment of ejection and the moment the droplet hits the well. Note this is equivalent between the I24 and NCI setup. C) A TTL signal is provided 1.2 ms after the XFEL pulse (red), signalling the motion stages to initiate movement (9–13 ms, depending on acceleration settings for the motorised stages). Subsequently, a second TTL signal signals the microdrop ejector to eject a droplet, which arrives 1–2 ms later on the fixed target chip (depending on ejector settings) after ejection. For this example, an arbitrary 13.5 ms incubation time was chosen.

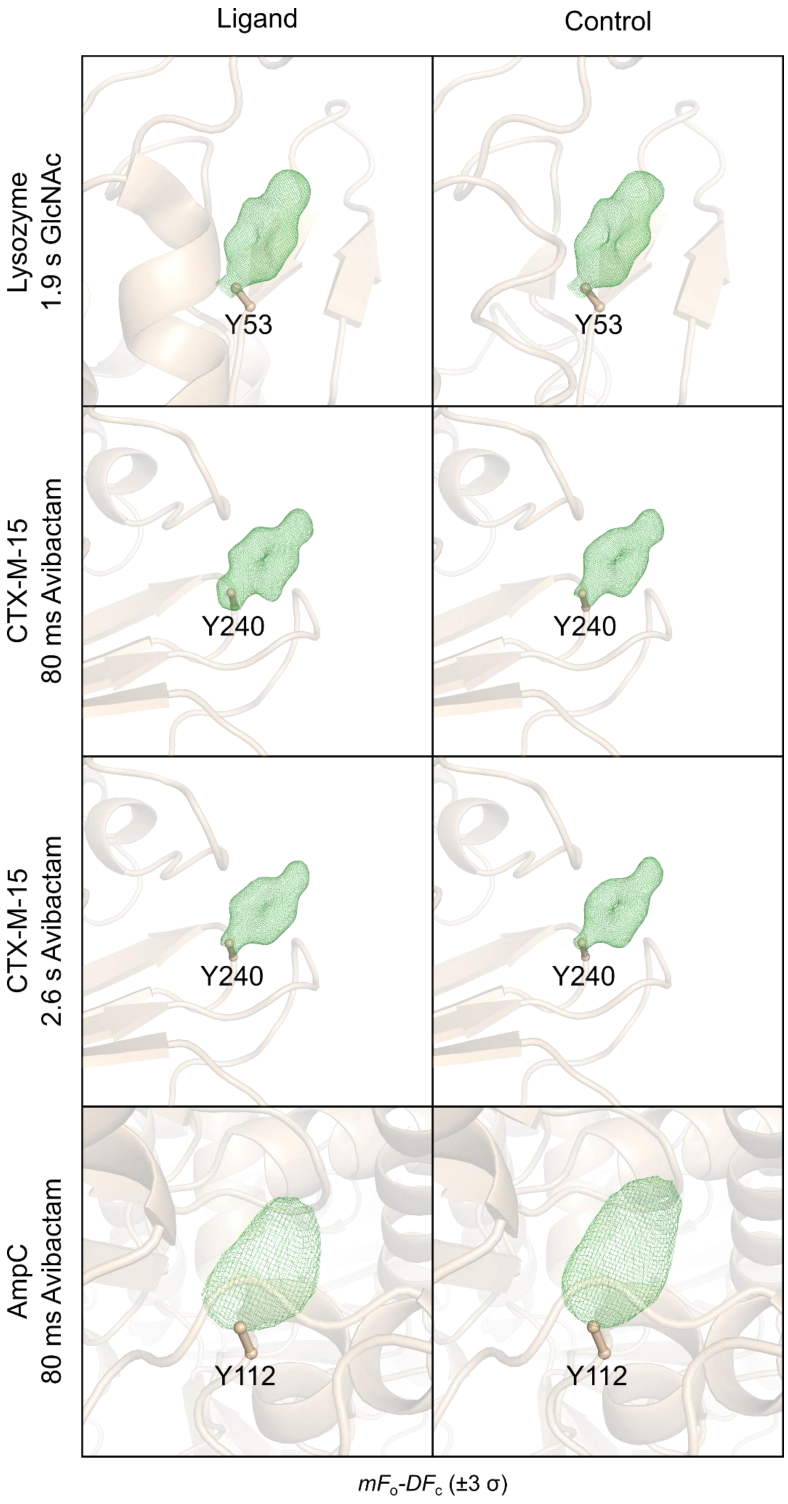

**Figure S9: Verification of data quality through OMIT maps of Tyr residues in the different enzymes.** $F_O$-$F_C$ maps (green, contoured at 3.0 σ) calculated after removal of the Tyr side chain by mutation to Ala followed by refinement. Residues selected were; Lysozyme Y53, CTX-M-15 Y240, $AmpC_{EC}$ Y112.

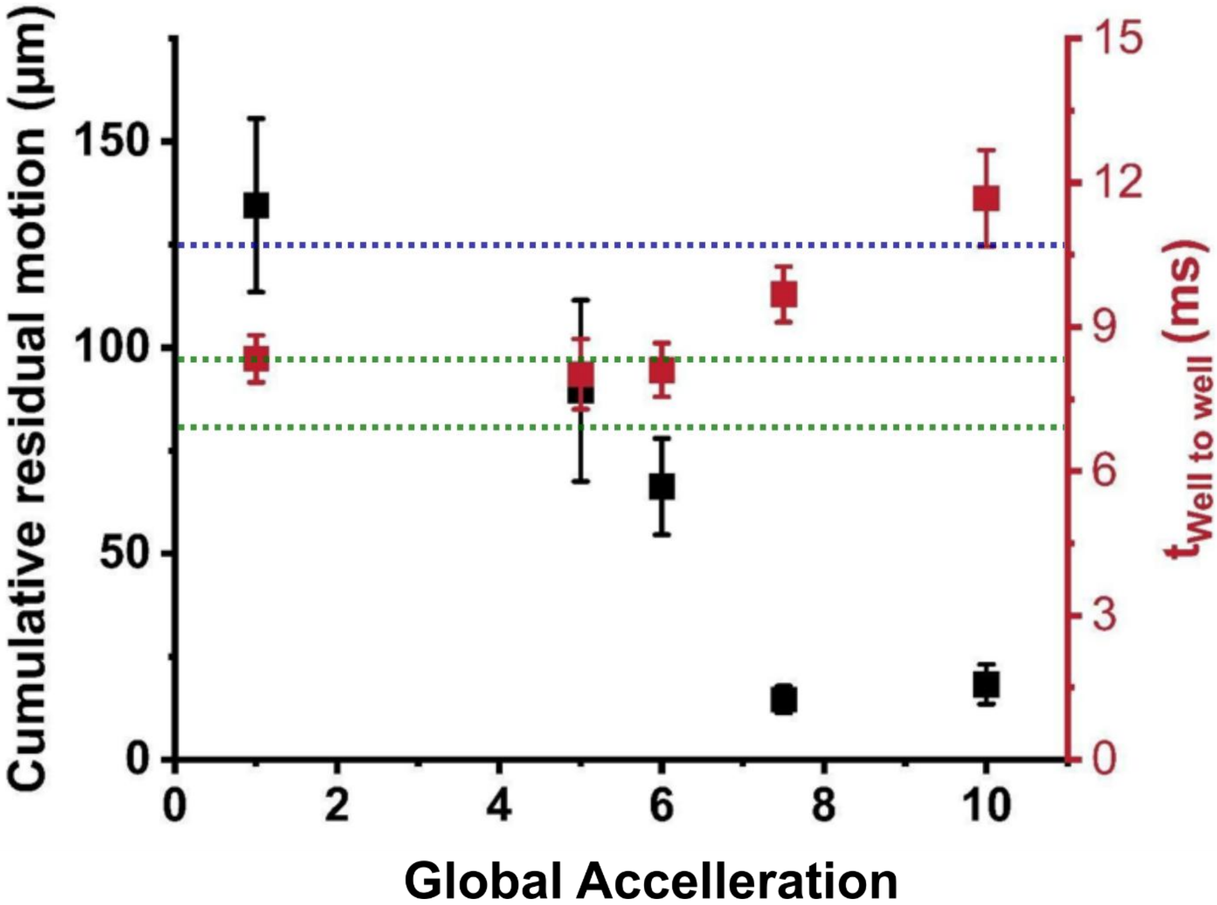

**Figure S10: Relation between global acceleration with cumulative residual motion and total travel time between adjacent wells within a row.** Global acceleration (GA) is a parameter set virtually to change acceleration and deceleration speeds. Stages were operated at a 30 Hz operation frequency; moving to 30 wells per s. Cumulative residual motion is the total amount of additional motion observed after arrival at the subsequent well averaged for >40 wells. Error bars are standard deviation on the observed distance per well. The blue line indicates inter well distance (125 µm), while the two green lines (98 and 83 µm) indicate the size of the open side of the chip with aperture size 20 and 5 µm, respectively. See Figure S1 and table S1 for further details.

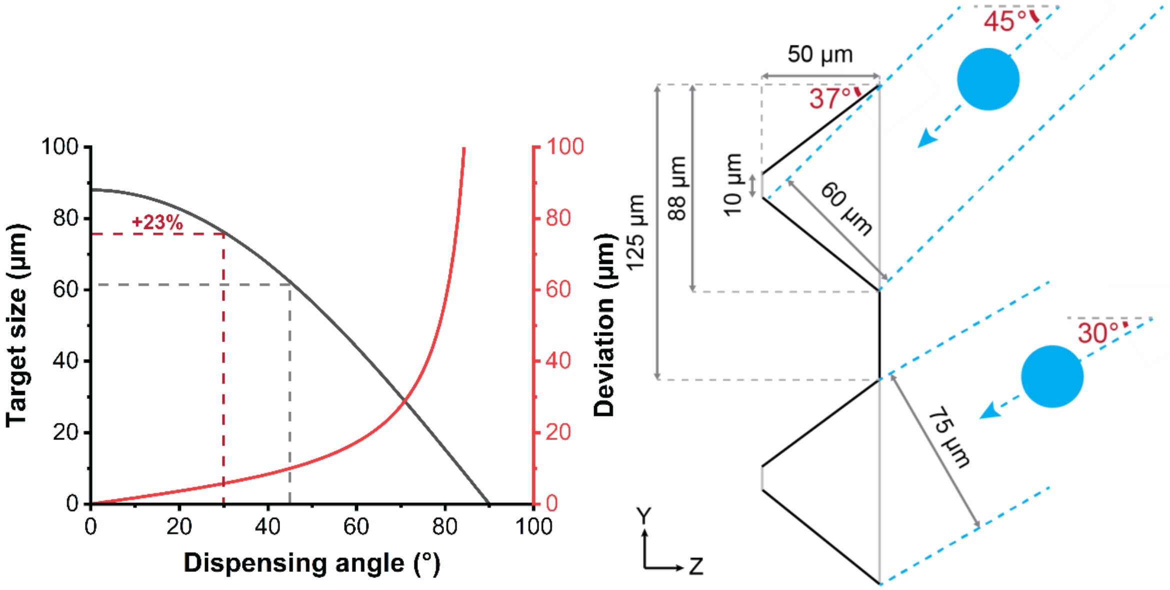

**Figure S11: Relationship between dispensing angle and target size deviation for a chip with 10 µm aperture wells.** Dispensing angle refers to the angle between the beam and the dispenser. Deviation refers to the distance between the well and the actual target hit, with an assumed displacement of a 5 µm in z position (derived from high speed camera footage) of the silicon chip as result of residual motion. The bottom part of the image is drawn to scale. Note the overall increase in target size (23%) when moving from 45° to 30° (*i.e.* 60° with respect chip plane) and the smaller deviation.

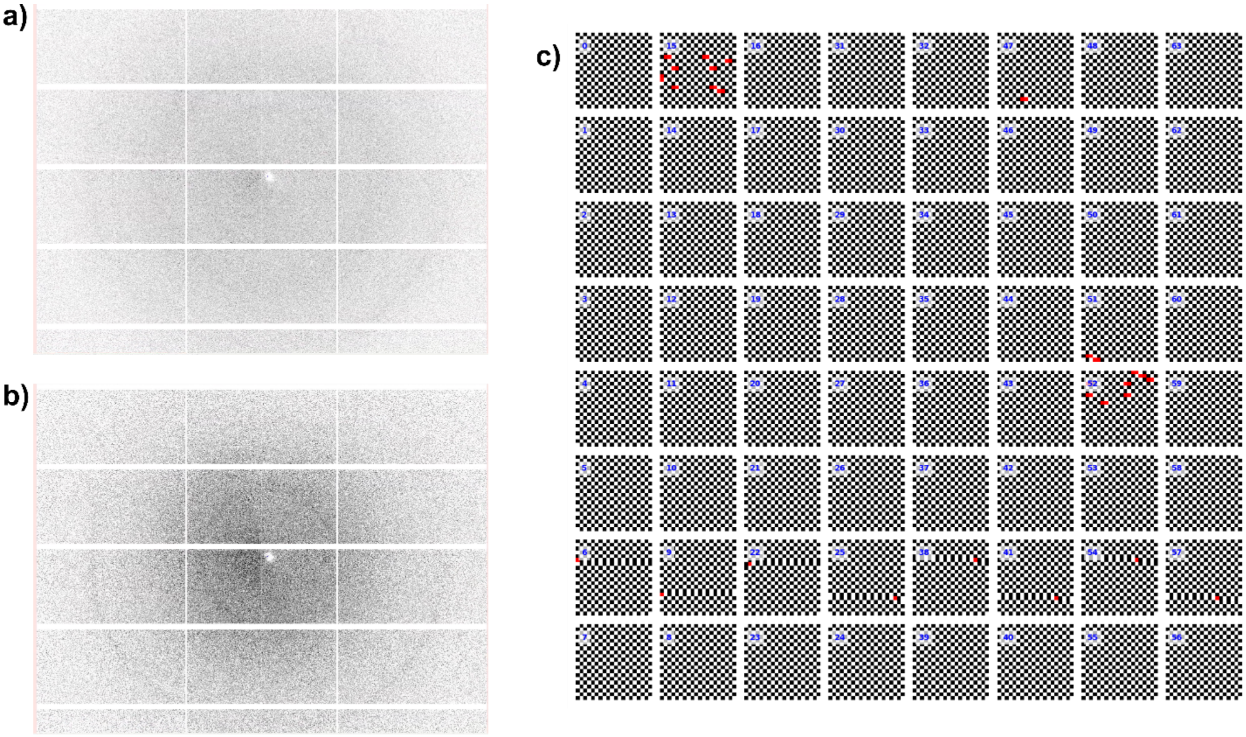

**Figure S12: Evaluation of droplet dispensing accuracy by water scattering on "dry" chip.** Water droplet dispensed in a checkerboard pattern on a dry silicon chip were hit with X-rays, causing radial scattering patterns for wells with droplets added (a) and blanc images (b) when no liquid was added. Total intensity of the image was used to evaluate droplet addition and no droplet addition and subsequently plotted (c). Black squares represent water add, while white squares represent empty wells, and red squares represent inaccurate ejection (*i.e.* water added where it should not or no liquid added where liquid was expected) (c). This example shows 25,600 wells of which 48 (0.188%) inaccurate droplet ejections.

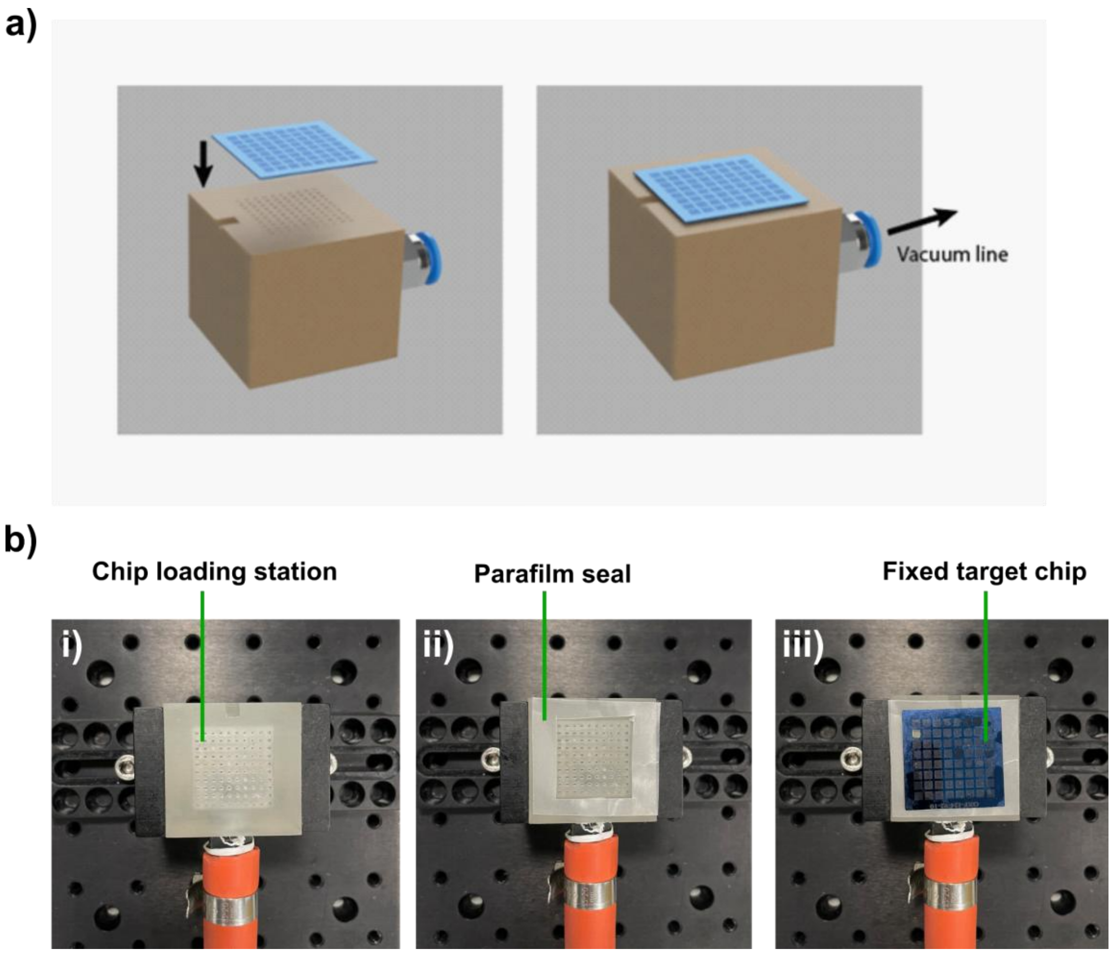

**Figure S13: Loading setup with loading station**. a) A 3D rendered image of the loading station, where a silicon chip can be loaded; b) photographs of the loading station during preparation of a fixed target chip. The loading station is housed within a humidity controlled enclosure and connected to a vacuum pump to facilitate removal of excess liquid. The chip can be placed directly on the custom 3D printed loading station or positioned on a parafilm seal (ii) enhancing the vacuum seal, improving efficient removal of liquid during the loading process.

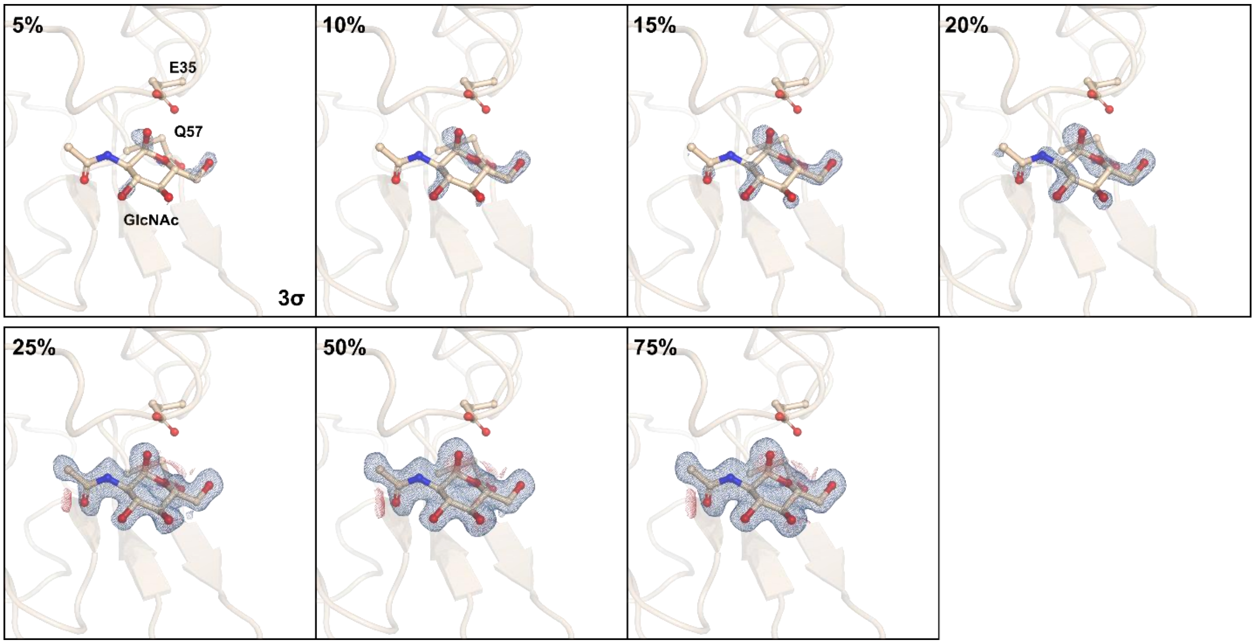

**Figure S14: Difference electron density maps between a partial N-acetyl-D-glucosamine (GlcNAc) bound Hen Egg White Lysozyme (HEWL) complex and apo-HEWL state.** Different percentages of HEWL•GlcNAc complex data mixed with apo-HEWL data, with a total of 10,000 reflections, are shown to demonstrate different levels of contamination of a control dataset. Data are determined to 1.7 Å, and derived from apo-HEWL and premixed HEWL•GlcNAc datasets. $F_o^{Partial\ complex}$-$F_o^{Apo}$ isomorphous difference map contoured at 3 σ, displaying the difference between the GlcNAc premixed HEWL and the ground state apo HEWL data. Note, a clear difference feature can be confidently determined at 5 and 10%, suggesting the sensitivity limit of this approach is around 5-10%.

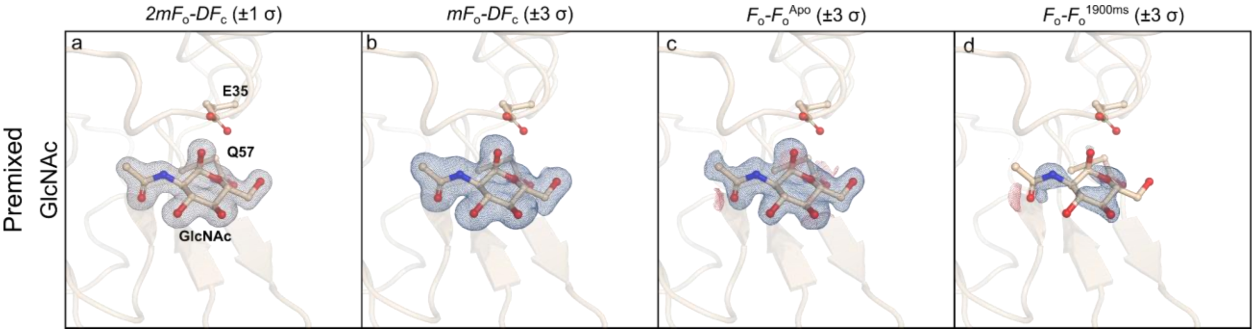

**Figure S15: Electron density map analyses of N-acetyl-D-glucosamine (GlcNAc) premixed Hen Egg White Lysozyme (HEWL) microcrystals obtained through established fixed target sample delivery.** Maps for HEWL mixed with GlcNAc (226 mM) for 10 minutes (a - d), collected using standard data collection approach, are determined to 1.64 Å resolution and were acquired at I24 Diamond Light Source. a) $2mF_o$-$DF_c$ electron density maps showing the active site of the HEWL•GlcNAc complex carved 1.5 Å around the ligand, contoured at 1 σ; b) $mF_o$-$DF_c$ polder OMIT difference map of the HEWL•GlcNAc complex carved 1.5 Å around the ligands and contoured at 3 σ; c) $F_o^{Premixed}$-$F_o^{Apo}$ isomorphous difference map contoured at 3 σ, displaying the difference between the GlcNAc premixed HEWL and the ground state apo HEWL data; d) $F_o^{premixed}$-$F_o^{1.9s}$ isomorphous difference map contoured at 3 σ, displaying the difference between the GlcNAc premixed HEWL and the GlcNAc added time point (1.9 s) data.

**Table S2:** Data collection and refinement statistics for mixing of GlcNAc with lysozyme (HEWL) recorded at I24, DLS

| | Resting state | 1.9 s mixing | 1.9 s control | Premixed |
|---|---|---|---|---|
| PDB ID: | 9TOQ | 9TOR | 9TOS | 9TOT |
| Wavelenght, (Å) | 0.9998 | 0.9998 | 0.9998 | 0.9998 |
| Space group | $P4_32_12$ | $P4_32_12$ | $P4_32_12$ | $P4_32_12$ |
| Unit cell | | | | |
| *a, b, c* (Å) | 79.23, 79.23, 38.22 | 78.98, 78.98, 38.29 | 78.96, 78.96, 38.26 | 79.16, 79.16, 38.11 |
| α, β, γ (°) | 90, 90, 90 | 90, 90, 90 | 90, 90, 90 | 90, 90, 90 |
| No. molecules / ASU | 1 | 1 | 1 | 1 |
| No. lattices | 25243 | 6781 | 6557 | 33400 |
| Resolution (Å) | 56.03–1.65 (1.78–1.65)* | 55.85-1.69 (1.82-1.69) | 55.83–1.67 (1.8–1.67) | 55.98-1.64 (1.77-1.64) |
| $R_{split}$ (%) | 5.2 (52.5) | 11.7 (73.3) | 12.7 (77.7) | 4.0 (32.3) |
| I/σI | 5.258 (0.362) | 2.308 (1.004) | 2.342 (0.795) | 7.597 (0.835) |
| $CC_{1/2}$ | 0.9965 (0.5364) | 0.9882 (0.3382) | 0.9840 (0.3420) | 0.9980 (0.5077) |
| Completeness (%) | 100 (100) | 100 (100) | 99.99 (99.86) | 100 (100) |
| Multiplicity | 270.4 (133.8) | 65.9 (33.1) | 48.18 (22.96) | 434.6 (203.2) |
| Wilson B value (Å$^2$) | 18.2 | 22.16 | 26.4 | 23.63 |
| ***Refinement*** | | | | |
| $R_{work}/R_{free}$ | 17.59/20.57 | 17.97/21.35 | 17.84/21.62 | 15.79/18.96 |
| No. atoms | | | | |
| Enzyme | 1053 | 1053 | 1049 | 1078 |
| Ligand | N/A | 20 | N/A | 20 |
| Water | 85 | 85 | 96 | 91 |
| Average B-factor | | | | |
| Enzyme | 27.31 | 25.44 | 25.41 | 26.8 |
| Ligands | N/A | 32.3 | N/A | 28.66 |
| Solvent | 38.85 | 37.21 | 37.12 | 39.32 |
| RMS deviations | | | | |
| Bond lengths (Å) | 0.005 | 0.006 | 0.006 | 0.006 |
| Bond angles (°) | 0.81 | 0.85 | 0.76 | 0.83 |

* highest resolution shell in parentheses, Detector: Pilatus3 6M

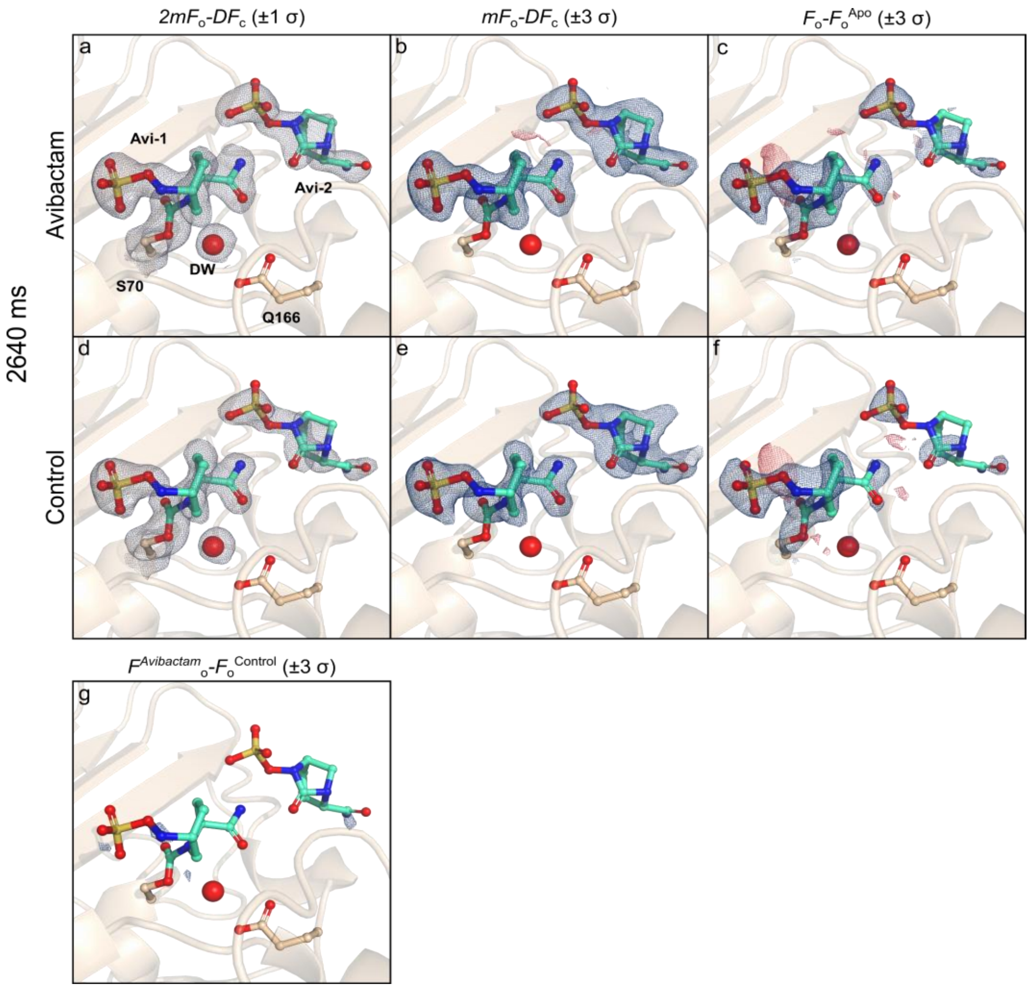

**Figure S16: Electron density map analyses of Avibactam (Avi) mixing with CTX-M-15 microcrystals obtained through drop on fixed target sample delivery demonstrating a high degree of cross well contamination.** CTX-M-15 mixed with Avi (250 mM) for 2.6 s (a - g), collected using Add then Revisit approach, are determined to 1.8 Å resolution and were acquired at I24 Diamond Light Source. a) 2mFo-DFc electron density maps showing the active site of the CTX-M-15•Avi complex carved 1.5 Å around relevant residues, contoured at 1 σ; b) $mF_o$-$DF_c$ polder OMIT difference map of the CTX-M-15•Avi complex carved 2.0 Å around the ligands and contoured at 3 σ; c) $F_o^{2.6s}$-$F_o^{Apo}$ isomorphous difference map contoured at 3 σ, displaying the difference between the Avibactam added time point (2.6 s) and the ground state apo CTX-M-15 data; d) $2mF_o$-$DF_c$ electron density map of the interleaved control (*i.e.* without ligand added) carved 1.5 Å around relevant residues, contoured at 1 σ; e) $mF_o$-$DF_c$ polder OMIT difference map of the interleaved control carved 2.0 Å around the ligands and contoured at 3 σ; f) $F_o^{Control}$-$F_o^{Apo}$ isomorphous difference map contoured at 3 σ, displaying the difference between the control (without ligand added) and apo CTX-M-15 data; g) $F_o^{2.6s}$-$F_o^{Control}$ isomorphous difference map contoured at 3 σ, displaying the difference between the avibactam added time point (2.6 s) and the control (without ligand added). Note the presence of electron density consistent with the presence of avibactam in the control $mF_o$-$DF_c$ polder OMIT (e) and isomorphous difference maps (f). In addition, the lack of difference features in the isomorphous difference map between the time point and control (g) further confirm cross well contamination.

**Table S3:** Data collection and refinement statistics for mixing of avibactam with CTX-M15 recorded at I24, DLS

| Time point<br>PDB ID | Resting<br>9TO5 | 2.6 s<br>9TOM | 2.6 s control<br>9TON | Contaminated time point<br>9TOO | Contaminated control<br>9TOP |
|---|---|---|---|---|---|
| ***Data collection*** | | | | | |
| Beamline (Wavelenght, Å) | 0.9998 | 0.9998 | 0.9998 | 0.9998 | 0.9998 |
| Space group | $P2_12_12_1$ | $P2_12_12_1$ | $P2_12_12_1$ | $P2_12_12_1$ | $P2_12_12_1$ |
| Unit cell | | | | | |
| *a*, *b*, *c* (Å) | 45.29, 45.94, 118.45 | 45.17, 45.86, 118.65 | 45.22, 45.71, 118.43 | 45.18, 45.82, 118.65 | 45.17, 45.80, 118.62 |
| α, β, γ (°) | 90, 90, 90 | 90, 90, 90 | 90, 90, 90 | 90, 90, 90 | 90, 90, 90 |
| No. molecules / ASU | 1 | 1 | 1 | 1 | 1 |
| No. lattices | 10756 | 9092 | 9152 | 3263 | 3058 |
| Resolution (Å) | 59.18–1.73 (1.65–1.6)* | 59.32 - 1.65 (1.7 - 1.65)* | 59.22–1.645 (1.7–1.65)* | 59.32–1.8 (1.88–1.8)* | 59.31–1.8 (1.88–1.8)* |
| $R_{split}$ (%) | 17.4 (45.3)* | 15.3 (63.8) | 15.3 (63.8)* | 30.8 (79.1)* | 30.0 (84.1)* |
| I/σI | 3.547 (1.341)* | 3.818 (1.075) | 3.373 (0.969)* | 2.173 (0.829)* | 2.179 (0.842)* |
| $CC_{1/2}$ | 0.956 (0.594)* | 0.9723 (0.3152) | 0.9564 (0.3088)* | 0.7961 (0.2874)* | 0.8650 (0.2855)* |
| Completeness (%) | 99.99 (99.89)* | 100 (100) | 100 (100)* | 99.39 (98.62)* | 99.41 (98.37)* |
| Multiplicity | 34.42 (9.18)* | 42.50 (16.60) | 40.23 (15.60)* | 14.58 (8.84)* | 13.45 (8.19)* |
| Wilson B value ($Å^2$) | 15.12 | 17.18 | 17.71 | 18.60 | 18.92 |
| ***Refinement*** | | | | | |
| $R_{work}/R_{free}$ | 14.06/16.61 | 0.1545/0.1885 | 17.57/20.50 | 0.1879/0.2299 | 0.1890/0.2296 |
| No. atoms | | | | | |
| Enzyme | 2012 | 2061 | 1994 | 2048 | 2061 |
| Ligand | N/A | 37 | N/A | 37 | 37 |
| Water | 218 | 196 | 201 | 214 | 200 |
| Average B-factor | | | | | |
| Enzyme | 18.53 | 19.9 | 20.18 | 20.65 | 21.38 |
| Ligands | N/A | 23.64 | N/A | 21.55 | 23.03 |
| Solvent | 32.75 | 33.88 | 32.31 | 32.58 | 33.52 |
| RMS deviations | | | | | |
| Bond lengths (Å) | 0.006 | 0.005 | 0.006 | 0.006 | 0.006 |
| Bond angles (°) | 0.85 | 0.77 | 0.77 | 0.8 | 0.8 |

* highest resolution shell in parentheses, Detector: Pilatus3 6M

**Table S4:** Data collection and refinement statistics for mixing of avibactam with CTX-M15 at NCI instrument, PAL-XFEL

| Time point | Resting | 80 ms | 80 ms control |
|---|---|---|---|
| PDB ID | 9TO1 | 9TOK | 9TOL |
| ***Data collection*** | | | |
| Wavelenght, (Å) | 1.30419 | 1.30420 | 1.30419 |
| Space group | $P2_12_12_1$ | $P2_12_12_1$ | $P2_12_12_1$ |
| Unit cell | | | |
| *a*, *b*, *c* (Å) | 45.29, 45.94, 118.45 | 45.28, 45.96, 118.49 | 45.28, 45.96, 118.49 |
| α, β, γ (°) | 90, 90, 90 | 90, 90, 90 | 90, 90, 90 |
| No. molecules / ASU | 1 | 1 | 1 |
| No. lattices | 16444 | 7681 | 5587 |
| Resolution (Å) | 42.83–1.60 (1.6276–1.60)* | 59.32–1.5 (1.54–1.5)* | 59.25–1.55 (1.59–1.55)* |
| $R_{split}$ (%) | 14.9 (38.3)* | 15.6 (75.7)* | 24.0 (68.0)* |
| I/σI | 4.286 (1.044)* | 3.569 (0.759)* | 2.682 (0.929)* |
| $CC_{1/2}$ | 0.9691 (0.6890)* | 0.9724 (0.3948)* | 0.9267 (0.3771)* |
| Completeness (%) | 99.99 (99.94)* | 100 (100)* | 99.99 (99.94)* |
| Multiplicity | 78.89 (11.77)* | 52.86 (18.56)* | 26.47 (14.39)* |
| Wilson B value (Å$^2$) | 15.12 | 17.59 | 18.00 |
| ***Refinement*** | | | |
| $R_{work}/R_{free}$ | 14.06/16.61 | 15.93/19.50 | 18.32/20.29 |
| No. atoms | | | |
| Enzyme | 2012 | 2015 | 2002 |
| Ligand | N/A | 24 | N/A |
| Water | 218 | 197 | 204 |
| Average B-factor | | | |
| Enzyme | 18.5 | 22.63 | 21.3 |
| Ligands | N/A | 32.82 | N/A |
| Solvent | 32.7 | 36.3 | 34.48 |
| RMS deviations | | | |
| Bond lengths (Å) | 0.006 | 0.006 | 0.005 |
| Bond angles (°) | 0.85 | 0.86 | 0.80 |

* highest resolution shell in parentheses, Detector: Rayonix MX225 HS

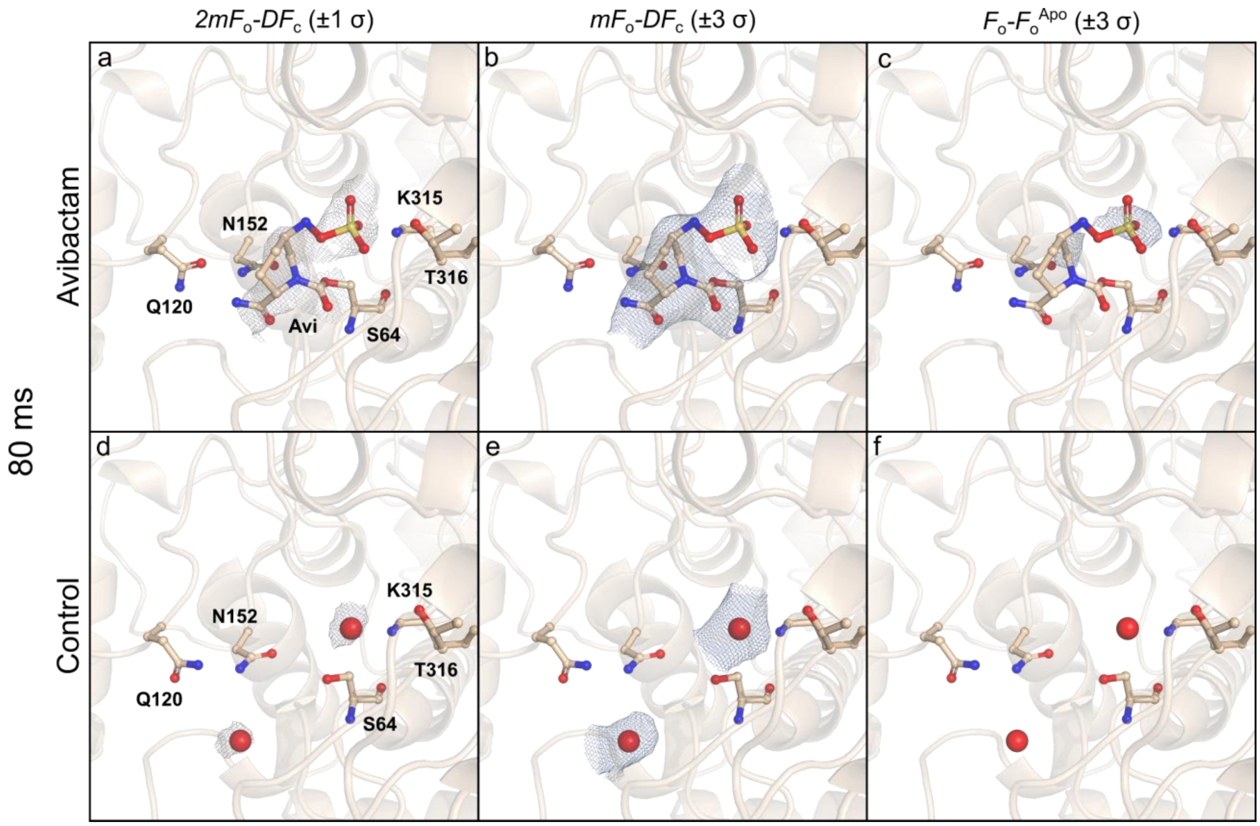

**Figure S17: Time resolved electron density map analyses of AmpC$_{EC}$ + Avibactam + 80 ms obtained through drop on fixed target sample delivery.** Maps for AmpC$_{EC}$ mixed with Avibactam (500 mM) and probed with X-rays after 80 ms (a - f) with the Add and Collect approach, were determined to 3.5 Å resolution and were collected at NCI PAL-XFEL. a) $2mF_o$-$DF_c$ maps of the active site of the partially occupied AmpC$_{EC}$•avibactam complex (radius: 1.5 Å, contour: 1 σ; b) $mF_o$-$DF_c$ polder OMIT map (radius: 1.5 Å, contour: 3 σ); c) $F_o^{80ms}$-$F_o^{Apo}$ isomorphous difference map (contour: 3 σ; integrated density: 8.71 e$^-$ Å$^{-3}$); d) $2mF_o$-$DF_c$ maps of the interleaved control (*i.e.* without ligand added) (radius: 1.5 Å, contour: 1 σ); e) $mF_o$-$DF_c$ polder OMIT map (radius: 1.5 Å, contour: 3 σ); f) $F_o^{Control}$-$F_o^{Apo}$ isomorphous difference map (contour: 3 σ; integrated density: 0.49 e$^-$ Å$^{-3}$).

**Table S5:** Data collection and refinement statistics for mixing of avibactam with $AmpC_{EC}$ at NCI instrument, PAL-XFEL

| Time point | Resting | 80 ms | 80 ms control |
|---|---|---|---|
| PDB ID | 9TOU | 9TOV | 9TOW |
| ***Data collection*** | | | |
| Wavelenght, (Å) | 1.30420 | 1.30420 | 1.30420 |
| Space group | *C*121 | *C*121 | *C*121 |
| Unit cell | | | |
| *a*, *b*, *c* (Å) | 118.71, 77.65, 98.75 | 119.07, 77.84, 99.31 | 118.71, 77.65, 98.75 |
| α, β, γ (°) | 90, 116.08, 90 | 90, 116.45, 90 | 90, 116.08, 90 |
| No. molecules / ASU | 2 | 2 | 2 |
| No. lattices | 5974 | 4777 | 5159 |
| Resolution (Å) | 45.25–1.88 (1.91–1.88) | 45.55–3.5 (3.85–3.5) | 45.57–2.78 (2.93–2.78) |
| $R_{split}$ (%) | 26.1 (99.9) | 20.6 (38.3) | 22.4 (43.4) |
| I/σI | 2.186 (0.539) | 3.818 (2.917) | 3.325 (1.609) |
| $CC_{1/2}$ | 0.9262 (0.0686) | 0.6826 (0.1315) | 0.9169 (0.3810) |
| Completeness (%) | 99.99 (99.94) | 99.97 (100) | 99.98 (99.90) |
| Multiplicity | 25.52 (17.34) | 37.95 (25.75) | 33.03 (21.70) |
| Wilson B value (Å$^2$) | 26.16 | 43.93 | 41.48 |
| ***Refinement*** | | | |
| $R_{work}/R_{free}$ | 18.38/22.24 | 13.22/21.98 | 15.88/21.70 |
| No. atoms | | | |
| Enzyme | 5598 | 5584 | 5606 |
| Ligand | N/A | 17 | N/A |
| Water | 450 | 37 | 103 |
| Average B-factor | | | |
| Enzyme | 28.94 | 27.81 | 34.99 |
| Ligands | N/A | 35.5 | N/A |
| Solvent | 37.77 | 23.13 | 36.76 |
| RMS deviations | | | |
| Bond lengths (Å) | 0.007 | 0.012 | 0.009 |
| Bond angles (°) | 0.84 | 1.25 | 1.00 |

Detector: Rayonix MX225 HS

### Methods

### Sample preparation

#### Hen Egg White Lysozyme (HEWL):

Microcrystals for Hen egg white lysozyme (HEWL) were prepared in batch using rapid-mixing as reported (Davy et al., 2019). In brief; a solution of HEWL (50 mg $mL^{-1}$; Sigma Aldrich L4919) in sodium acetate (20 mM, pH 4.6) was mixed through vortexing with an equal volume of crystallisation solution (citric acid (1 M), NaCl (20 w/v%), PEG 6,000 (5 w/v%), pH 3.0) in a polypropylene tube (1.5 mL) at 22 ºC. Nucleation of the microcrystals was near instant and the resulting suspension was left for 1 h to mature the crystals (15–20 µm; ~$10^8$ crystals $mL^{-1}$). Crystal sizes were determined using a Hirox HRX-01 microscope (20 to 5000 magnification), while crystal density was determined using a Neubauer counting chamber.

#### CTX-M-15:

Recombinant His-tagged CTX-M-15 was expressed, purified and crystallised as previously described (Butryn et al., 2021). In brief, His-tagged CTX-M-15 (pOPINF expression vector) was recombinantly expressed in SoluBL21 (DE3) E. coli and purified using Ni-NTA resin (Qiagen; preequilibrated in HEPES (50 mM, pH 7.5) and NaCl (400 mM)) and eluted with imidazole (400 mM). The His-tag was removed with 3C protease (4 °C, overnight) and binding to incubation with Ni-NTA resin. After size exclusion chromatography (Superdex 75; preequilibrated with 50 mM HEPES pH 7.5, 150 mM NaCl ) the desired fractions were pooled and concentrated (20 mg $mL^{-1}$) by centrifugation. Microcrystals of CTX-M-15 were grown in 24-well sitting drop plates (Hampton) by mixing protein (5 µL) with crystallisation solution (5 µL; 2.0 M $(NH4)_2SO_4$, 0.1 M Tris pH 8) and of seeds (5 µL; generated by crushing macro CTX-M-15 crystals), and equilibrating against crystallisation solution (500 µL). Crystals grew as rods within 24 h, with a maximum width of 5 µm and length of 10–20 µm. Crystal density was ~$1 \cdot 10^8$ crystals $mL^{-1}$ as measured using a TC20 automated cell counter (BioRad) and Neubauer counting chamber.

#### $AmpC_{EC}$:

Recombinant $AmpC_{EC}$ from *Escherichia coli* was prepared using a modified version of a reported protocol (Page, 1993; Lang *et al.*, 2020). In brief, $AmpC_{EC}$ (pAD7 expression vector; tetracycline) was expressed in E. coli W3110 cells using 2TY media. Cells were grown overnight (37 °C) and harvested by centrifugation (10 min, 12,000 ×g, 4 °C). After resuspending in lysis buffer (Tris (20 mM, pH 7.5), $MgCl_2$ (10 mM), DNAse I (50 µg $mL^{-1}$)) the cells were lysed through sonification and debris removed by centrifugation. The supernatant was dialysed against dialysis buffer (Tris (10 mM, pH 6.75), thrice), before cation exchange purification (SP Sepharose (GE Healthcare, Chicago, USA) and eluted using a gradient (Tris, 10–75 mM, pH 7.0). Fractions containing $AmpC_{EC}$ were pooled and concentrated to 25 mg $mL^{-1}$ using a 10 kDa molecular weight cutoff centrifugal concentrator. Initial crystals of $AmpC_{EC}$ were obtained through hanging drop vapour diffusion; $AmpC_{EC}$ (1.5 µL) was added to precipitant solution (4.5 µL, potassium phosphate (1.6 M, pH 8.5)) on unsiliconised coverslides, and equilibrated over precipitant solution (500 µL) at room temperature (1–2 weeks). Macro $AmpC_{EC}$ crystals were obtained and crushed using a seed beat kit (Hampton Research, Aliso Viejo, USA); alternating between vortexing (30 s), followed by chilling on ice (30 s; 4 °C). The seed stock (diluted 10–10,000×) was used to prepare macro $AmpC_{EC}$ crystals on siliconised coverslides (3–4 days), which were then used to prepare a new microcrystal seed stock. The concentrated seed stock was used in batch to obtain $AmpC_{EC}$ microcrystal slurries; precipitant solution (900 µL, potassium phosphate (1.9 M, pH 8.8)) was mixed with $AmpC_{EC}$ (100 µL, 50 mg $mL^{-1}$), followed by the addition of microseed stock (50 µL). Slurries of $AmpC_{EC}$ microcrystals grew in 2–4 h (2.5 × 6 × 2 $µm^3$, 2.4 · $10^7$ crystals $mL^{-1}$) and used without further concentration.

**Optimisation of droplet dispensing parameters**

To operate the Autodrop Pipette (Microdrop Technologies) most effectively, various solutions were prepared and tested prior the experiment. Excess of small molecule solution (>500 µL), preferentially in an aqueous solution, was filtered and loaded into the pipette (~75 µL maximum capacity) according to guidelines provided in the manual. Although theoretically capable of loading DMSO solution, in general our experience showed more difficulty dispensing highly concentrated (>100 mM) small molecule solutions. Ejection stability was monitored using a camera at fixed frame rate and a stroboscopically operated LED light, over a distance of ~1 mm at while switching back and forth between different dispensing frequencies (1 Hz, 30 Hz), while periodically interrupting dispensing. It is important to note, that dispensing parameters were pipette bound; *i.e.* switching to a different Autodrop Pipette of the same specifications, would require some optimisation of the dispensing parameters; however, the previously found conditions, often provided a good starting point. The pipette was usually operated in “triple pulse mode”, which providing multi-pulse actuation waveform. This mode generated smaller volume droplets and allowed for a higher degree of control over ejection properties compared to the “single pulse mode”.

These tests were aimed at providing robust dispensing parameters

**Determination of droplet hit accuracy**

An empty silicon chip mounted in a chip holder was attached to the motorised stages via a kinematic mounting on I24, Diamond Light Source. Droplets of Milli-Q water (~90 pL) were dispensed using the PEI and data were collected using the "Add & Collect" method, with the shortest possible delay time (*i.e.* 10 ms acquisition).

Radial averaging of the scattering observed in the images were used to distinguish "hit" and "empty" wells. Deviations from the expected pattern were established using a python script and the number divided over the total number of wells to obtain the hit percentage.

**Generating synthetically contaminated data sets**

Data collected on resting (apo) state hen egg white lysozyme (HEWL) and *N*-acetyl-D-glucosamine (GlcNAc) premixed with HEWL, were indexed and integrated using DIALS. Scaling and merging of the data were done combining different amounts (*i.e.* 5-75%) of random reflections of the premixed state into the apo state to a total of 10,000 lattices, using cctb.xfel.merge (Hatte *et al.* 2014) and with a GlcNAc mixed lysozyme pdb file (ID: 7BHN) as reference. The resolution of the data was cut at 1.7 Å, molecular replacement done with phenix.phaser, followed by automated refinement using phenix.refine.

**Integrating isomorphous difference density maps**

Isomorphous difference density maps were generated using phenix.fobs_minus_fobs_map, contoured at 3 σ, with 1.5 Å radius. Integration of Fourier difference map features was achieved using in house python scripts. The σ-values in each map were extracted from mtz files using mapdump CCP4 package (Agirre *et al.* 2023). Using the difference structural factors, all maps were regenerated with identical sampling rate via the Gemmi crystallographic library (version 0.7.4; Wojdyr, 2022), and selecting the ligand of interest for integration. All positive and negative values above the 3 σ cut-off within 1 Å radius from the ligand were integrated.

**Diffusion experiments by NMR**

Compounds, including avibactam (supplier), were utilised in diffusion NMR spectroscopy (DOSY) experiments without additional purification. Stock solutions of avibactam were prepared in $D_2O$ (>99% $^2$H, Sigma-Aldrich) and diluted appropriately in $D_2O$ (>99% $^2$H). Diffusion experiments were conducted in 5 mm NMR tubes (Norell) at 298 K on a Bruker AVIII 500 equipped with a BBO probe. Data acquisition was performed using LED experiments with a bipolar gradient (ledbpgp2s) and a squared gradient ramp of 16 steps between 5 and 95%, a gradient pulse length of 1.5 ms and a diffusion time of 100 ms. Each step consisted of an 8-scan acquisition. Suitable non-overlapping peaks were selected for integration and fitting with equation (1) using Bruker Topspin 3.6.1. Calculated results for distinct peaks were averaged to determine the final diffusion coefficient.

$$I_G = I_{G=0} e^{-(\gamma \delta G)^2 D\left(\Delta - \frac{\delta}{3}\right)} \quad (1)$$

In which; $I$ = measured signal intensity, $I_G$=0 = signal intensity for $g$ = 0, $\gamma$ = magnetogyric ratio, $D$ = diffusion coefficient, $G$ = gradient strength, $\Delta$ = diffusion time, $\delta$ = gradient pulse length.

CTX-M-15 crystals display cuboid morphology with dimensions ranging from 3-8 by 10-20 µm. Given the morphology, the diffusion time calculation can be reduced to a 1D diffusion problem, looking at the shortest dimension. On average, the crystal dimensions are estimated to be 15 × 15 × 5 µm$^3$. Following Fick's 1$^{st}$ and 2$^{nd}$ law of diffusion (Schmidt, 2020):

$$D\nabla^2 C = \frac{\partial C}{\partial t}$$

in which $D$ = diffusion coefficient, and $\nabla^2$ = Laplace operator, $C$ = time and space dependent concentration, and $t$ = time. Applying crystal constraints of dimension a, b, and c (15, 15, 5 µm); and $D_{avibactam} = 5.65 \cdot 10^{-6}$ cm$^2$ s$^{-1}$, $N$ = 20, $C_0$ = ligand concentration outside the crystal, assuming a final [Avibactam] of 50% (*i.e.* $C_{\mathrm{centre}}(t_{50\%})/C_0 = 0.5$):

$$C(x,y,z,t) = C_0 \Bigg[1 - \frac{64}{\pi^3} \sum_{l=0}^{N} \sum_{m=0}^{N} \sum_{n=0}^{N} \frac{-1^{l+m+n}}{(2l+1)(2m+1)(2n+1)} \times \cos\frac{(2l+1)\pi x}{2a} \cos\frac{(2m+1)\pi y}{2b} \cos\frac{(2n+1)\pi z}{2c} \times exp\left[-D\frac{\pi^2}{4}\left(\frac{(2l+1)^2}{a^2} + \frac{(2m+1)^2}{b^2} + \frac{(2n+1)^2}{c^2}\right)\right]\Bigg]$$

this gives, $t_{50\%} \approx 4.18$ ms for avibactam with average CTX-M15 crystals.
